# A plasmid locus associated with *Klebsiella* clinical infections encodes a microbiome-dependent gut fitness factor

**DOI:** 10.1101/2020.02.26.963322

**Authors:** Jay Vornhagen, Christine M. Bassis, Srividya Ramakrishnan, Robert Hein, Sophia Mason, Yehudit Bergman, Nicole Sunshine, Yunfan Fan, Winston Timp, Michael C. Schatz, Vincent B. Young, Patricia J. Simner, Michael A. Bachman

## Abstract

Gut colonization by the pathogen *Klebsiella pneumoniae* (Kp) is consistently associated with subsequent Kp disease^1-5^, and patients are predominantly infected with their colonizing strain^1,2^. However, colonizing strains likely vary in their potential to cause infection. We previously identified the plasmid-encoded tellurium resistance (*ter*) operon as highly associated with infection when compared to asymptomatic colonization in hospitalized patients^1^. The *ter* operon bestows resistance to the toxic compound tellurite oxide (TeO_3_^−2^), but this is unlikely to be its physiological function, as tellurium and TeO_3_^−2^ are exceedingly rare. Here we show that *terC* is necessary and *terZABCDEF* is sufficient for phenotypic TeO_3_^−2^ resistance. Next, we demonstrate that *ter* is encoded on a diverse group of plasmids without known plasmid-encoded virulence genes, suggesting an independent role in infection. Finally, our studies indicate that *ter* is a gut fitness factor, and its fitness advantage is conferred only when specific gut microbiota constituents are present. Collectively, these data reveal the Kp *ter* operon that is highly associated with human infection likely acts early in pathogenesis as a horizontally-transferrable fitness factor promoting robust gut colonization in the presence of the indigenous microbiota.

---

Given the strong association between *ter* and disease-causing Kp^1^, we aimed to characterize this locus, whose biology has long been enigmatic. The antibacterial property of TeO_3_^−2^ was first described by Sir Alexander Fleming in 1932^6^, mediated through the reduction of TeO_3_^−2^ to Te^0^ in bacterial cells^7^. Given the rarity of Te^0^ and the fact that TeO_3_^−2^ is toxic in humans^8^, we postulated that resistance to TeO_3_^−2^ is a robust *in vitro* phenotype but not directly relevant to infection. Thus, we aimed to determine the impact of *ter in vivo*. Previous *in silico* studies show that the *ter* locus is present in many different bacterial families^9-11^. The complete locus consists of two distinct, but highly conserved, operons encoded on opposite DNA strands: a set of 14 individual genes with potential biosynthetic functionality that includes *terXYW*, and a tellurium resistance operon consisting of *terZABCDEF.* This is followed by a predicted haloacid dehydrogenase (HAD) that is highly conserved among *ter*-encoding Kp isolates (Fig. 1a, Supplementary Table 1). Deletion of *terC* from the pK2044 plasmid renders Kp strain NTUH-K2044 exquisitely sensitive to TeO_3_^−2^ (Fig. 1b). Computational prediction of protein structure and function based on amino acid sequence using I-TASSER^12-14^ indicates that *terC* is involved in transmembrane transport (Supplementary Table 1). *terC* expressed *in trans* in a Δ*terC* strain did not restore phenotypic resistance to TeO_3_^−2^ (Fig. 1b), attributable to insufficient plasmid expression or polar effects of the mutation. Instead, *terZ-F* expressed *in trans* fully restored TeO_3_^−2^ resistance (Fig. 1b). Furthermore, the expression of *terZ-F in trans* was sufficient to confer TeO_3_^−2^ resistance to the *Escherichia coli* strain MG1655 (Fig. 1c).

**Fig. 1:**
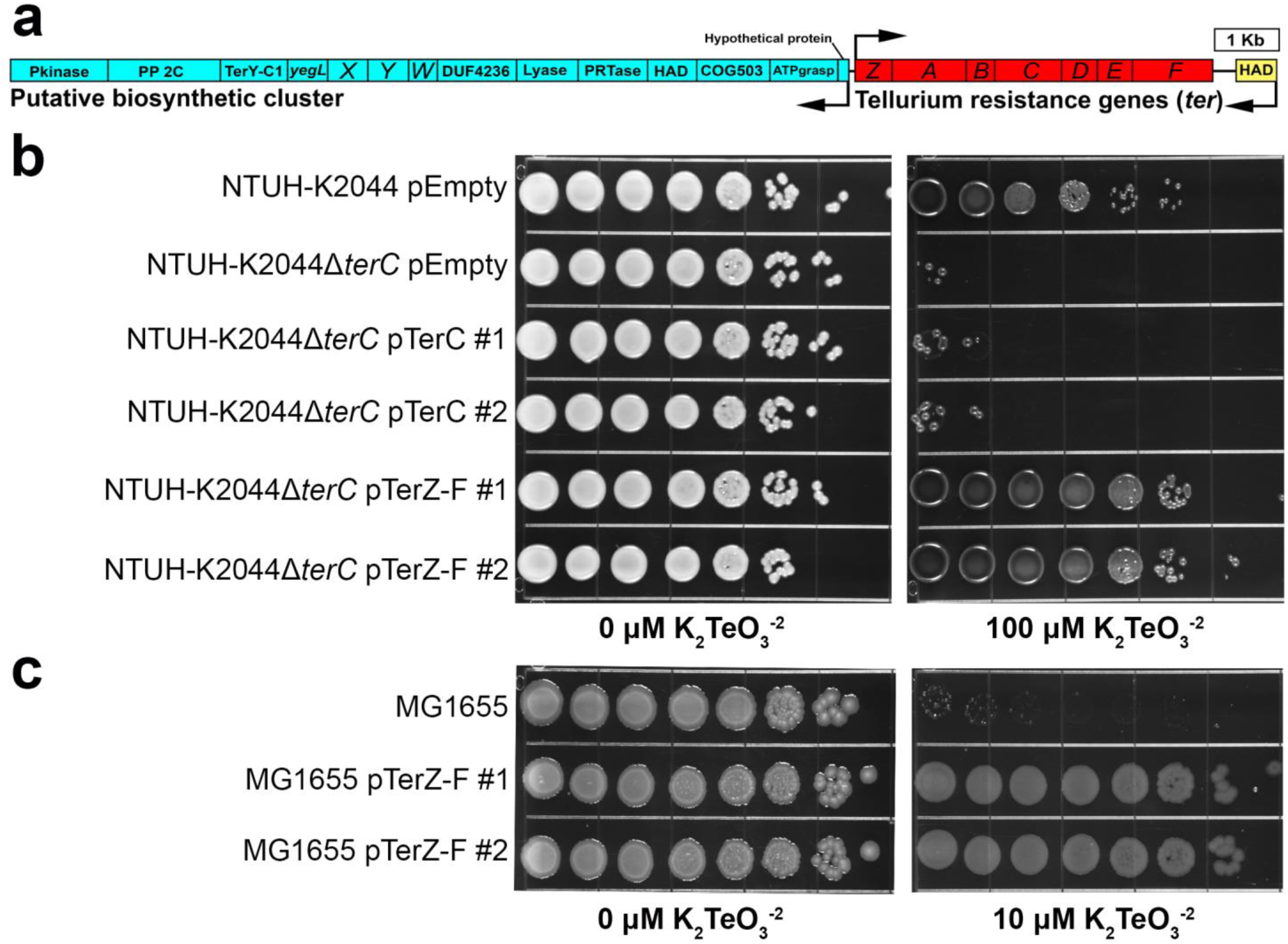
The Kp *terZ-F* genes are sufficient for TeO_3_^−2^ resistance. **a**, The *ter* locus is organized in two operons, a putative biosynthetic cluster and a tellurium resistance cluster. These sections are found on opposite DNA strands and are encoded in bidirectionally. The representative *ter* locus from the hvKp strain NTUH-K2044 is shown. **b, c**, NTUH-K2044 containing an empty vector, the isogenic Δ*terC* mutant containing an empty vector, the pTerC, or the pTerZ-F plasmid (**b**), and the *E. coli* K12 strain MG1655 with or without the pTerZ-F plasmid (**c**) were grown on LB or LB containing 10 or 100 μM potassium tellurite to visualize inhibition of growth. Two representative clones of NTUH-K2044Δ*terC* containing the pTerC or the pTerZ-F plasmid and MG1655 containing the pTerZ-F plasmid are shown.

Previous studies demonstrate that *ter* can be found on pK2044-like plasmids that encode multiple virulence genes characteristic of hypervirulent Kp strains (hvKp)^15,16^. This suggests the association between *ter* and Kp disease could be due to genetic linkage with plasmid-encoded virulence genes. To determine if this is the case, we characterized the *ter*-encoding (*ter*+) Kp isolates and plasmids from our previous study. These isolates were highly diverse^1^, as reflected by their *wzi* capsule types^17^ (Fig. 2a) and none were a hvKp type. To determine virulence gene content, we sequenced *ter*-encoding plasmids from our patient isolates using Nanopore technology. *ter*-encoding plasmids were characteristically large (86.8-430.8 kb; Supplementary Fig. 1) and some were derived from plasmid fusions (Supplementary Table 2). These plasmids displayed substantial variation in sequence similarity outside the *ter* locus (Supplementary Fig. 2a-b), indicating that *ter* is not a marker for a widely circulating, highly conserved, plasmid. Additionally, these *ter*-encoding plasmids do not contain virulence genes associated with Kp hypervirulence, nor was a single antibiotic resistance gene highly present on these plasmids (Fig. 2b, Supplementary Table 2, 3). This supports the premise that *ter* is an independent fitness factor during infection, rather than a marker of a hypervirulence or antibiotic resistance-encoding plasmid. Finally, predicted open reading frames (ORFs) directly up- and downstream of the *ter* operon displayed minimal functional conservation (Fig. 2c), further indicating that *ter* is not tightly linked to a virulence factor. Next, we repeated this analysis using publicly available genomes of *ter*-encoding Kp isolates (n = 88). These isolates were not limited to hvKp *wzi* types (Fig. 2d), and indeed, the *ter+* plasmids displayed variable sequence similarity (Supplementary Fig. 2c). 42% of *ter*+ plasmids contained a *rmpA/A2* homologue and an accessory iron acquisition system. The remaining 58% *ter*+ plasmids had no classical hypervirulence factor present (Fig. 2e, Supplementary Table 2, 3), and again, predicted up- and downstream ORFs displayed little functional conservation with the exception of transposase activity (Fig. 2f, Supplementary Table 1). Together, these results indicate that *ter* is a genetically independent factor, and is likely playing a direct, but yet undescribed role in Kp disease.

**Fig. 2:**
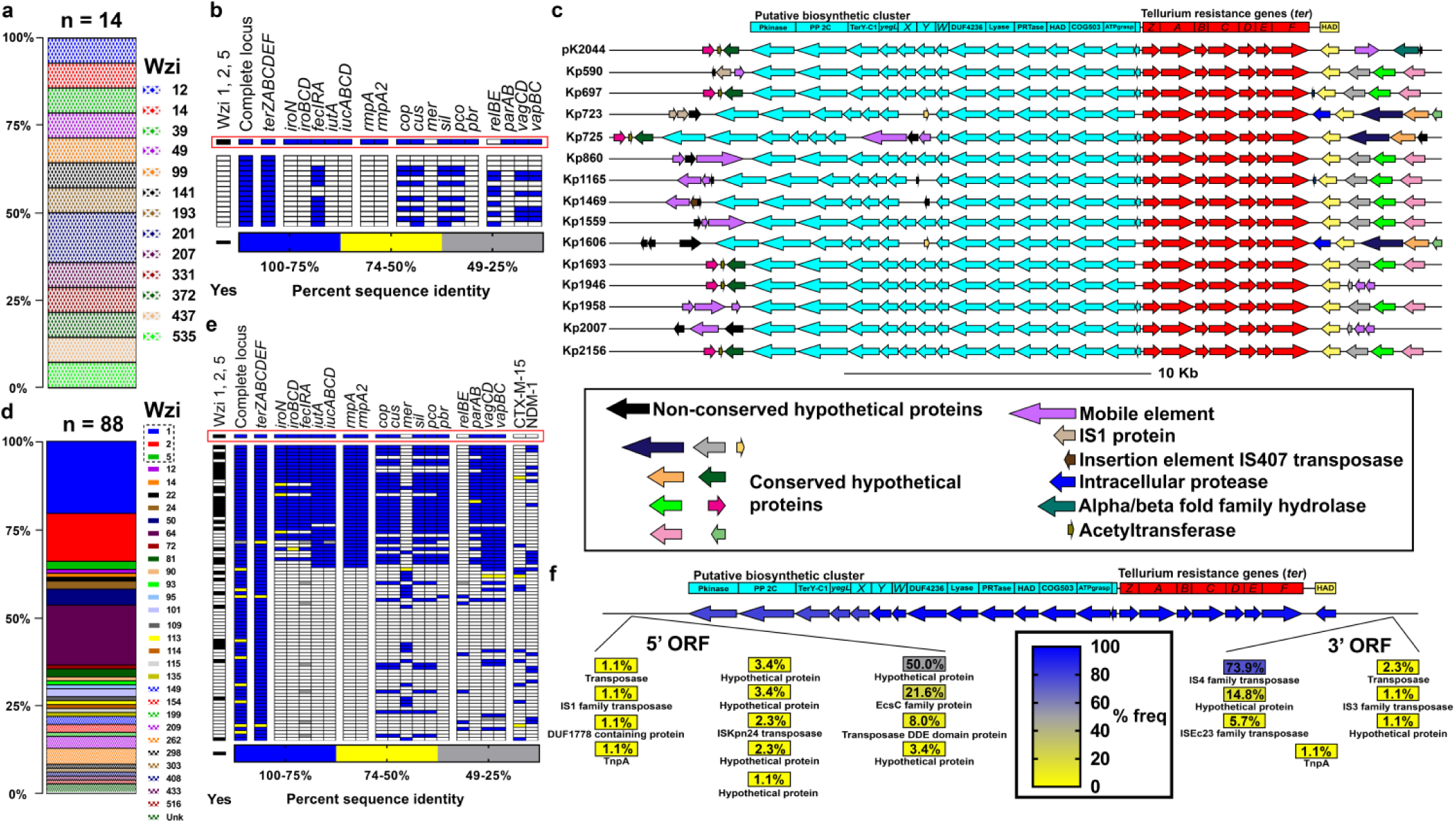
The Kp *ter* operon is not exclusive to hypervirulence plasmids. *ter*+ plasmids from Martin *et al.* mSystems, 2018^1^ (**a-c**) and reference strains from the NCBI database (**d-f**) were analyzed. **a**,**d**, Relative frequencies of *wzi* types of Kp strains containing *ter*+ plasmids. HvKp Wzi types are outlined in black. **b**,**e**, Heat map of *ter*+ plasmid sequence similarity to genes known to influence infection and antibiotic resistance genes. Each row represents an individual plasmid in the order of Supplementary Table 2 (Martin *et al.* mSystems, 2018^1^ index 1-14, NCBI reference strains index 15-102). The pK2044 hvKp plasmid is highlighted by the red box, and hvKp Wzi types are indicated. **c**,**f**, To determine if any neighboring gene was consistently associated with *ter*, the gene neighborhood of *ter* plasmids encoding the *ter* operon from Martin *et al.* mSystems, 2018^1^ was visualized (**c**) and the frequency of ORFs adjacent to the *ter* operon encoded on reference plasmids from the NCBI database was calculated (**f**).

We previously reported a strong association between *ter* and Kp infection (pneumonia and bacteremia) in colonized patients, yet also found that *terC* is dispensable in a murine model of pneumonia^1^. To determine if *ter* is important for bloodstream infection, WT Kp and Δ*terC* were competed in a peritoneal injection model of murine bacteremia. Δ*terC* was dispensable in all tissues with the exception of a modest defect the brain (Supplementary Fig. 3). We did not explore this finding further, as a role in meningitis would not explain the correlation between *ter* and infections observed in patients. We then hypothesized that *ter* may be required during gut colonization, which precedes infection^2,3^. Exposure to antibiotics was not associated with Kp colonization or subsequent infection in our intensive care unit patient population^1,18^, indicating that Kp must contend with the indigenous microbiota to colonize the gut and cause infection. Previous studies have shown that gut microbiota differs by mouse vendor and by the room within each vendor facility (referred to as “barrier”)^19^ and these differences can impact experimental output^20-22^. Therefore, C57BL/6J mice were sourced from two different barriers (RB16 and RB07) at The Jackson Laboratory and inoculated orally with a mixture of wild-type and Δ*terC* Kp (Fig. 3a). Intriguingly, a fitness defect (median 5.8, 4.7, 8.9, and 4.0 fold-defect on days 1-4, respectively) was observed consistently for Δ*terC* in the mice sourced from RB16 over several days (Fig. 3b, Supplementary Fig. 4a) but not from RB07, despite their genetic identity (Fig. 3c, Supplementary Fig. 4b). These data suggest that the fitness defect exhibited by Δ*terC* is dependent on the gut microbiota. To test this hypothesis, mice were treated with antibiotics to disrupt the indigenous microbiota, and the experiment was repeated. Disruption of the gut microbiota increased overall Kp colonization density in mice from both barriers (Supplementary Fig. 4c,d) and restored the fitness of Δ*terC* in mice sourced from RB16 (Fig. 3d, Supplementary Fig. 4c). Conversely, antibiotic treatment of RB07 mice did not impact Δ*terC* fitness (Fig. 3e, Supplementary Fig. 4d). These data indicate that the microbiota of barrier RB16 are involved in reducing the fitness of the Δ*terC* mutant, as opposed to the microbiota of RB07 enhancing Δ*terC* fitness. Expression of *terZ-F in trans* partially complemented the Δ*terC* fitness defect in RB16 mice (Fig. 3f). Next, we aimed to determine if *ter* is required for colonization by performing mono-colonization studies. These results show that both strains were able to colonize the gut (Supplementary Fig. 5), although there was significant mouse-to-mouse variation that may obfuscate an advantage of *ter* during mono-colonization. Consistent with previous studies^23-26^, the mice exhibited increased mortality throughout the duration of the experiment. However, the increased bacterial load due to antibiotics (Supplementary Fig. 4) was not associated with increased mortality, and both strains exhibited equal virulence in this model regardless of barrier (Supplementary Fig. 6). Collectively, these data suggest that *ter* is a microbiome-dependent gut fitness factor.

**Fig. 3:**
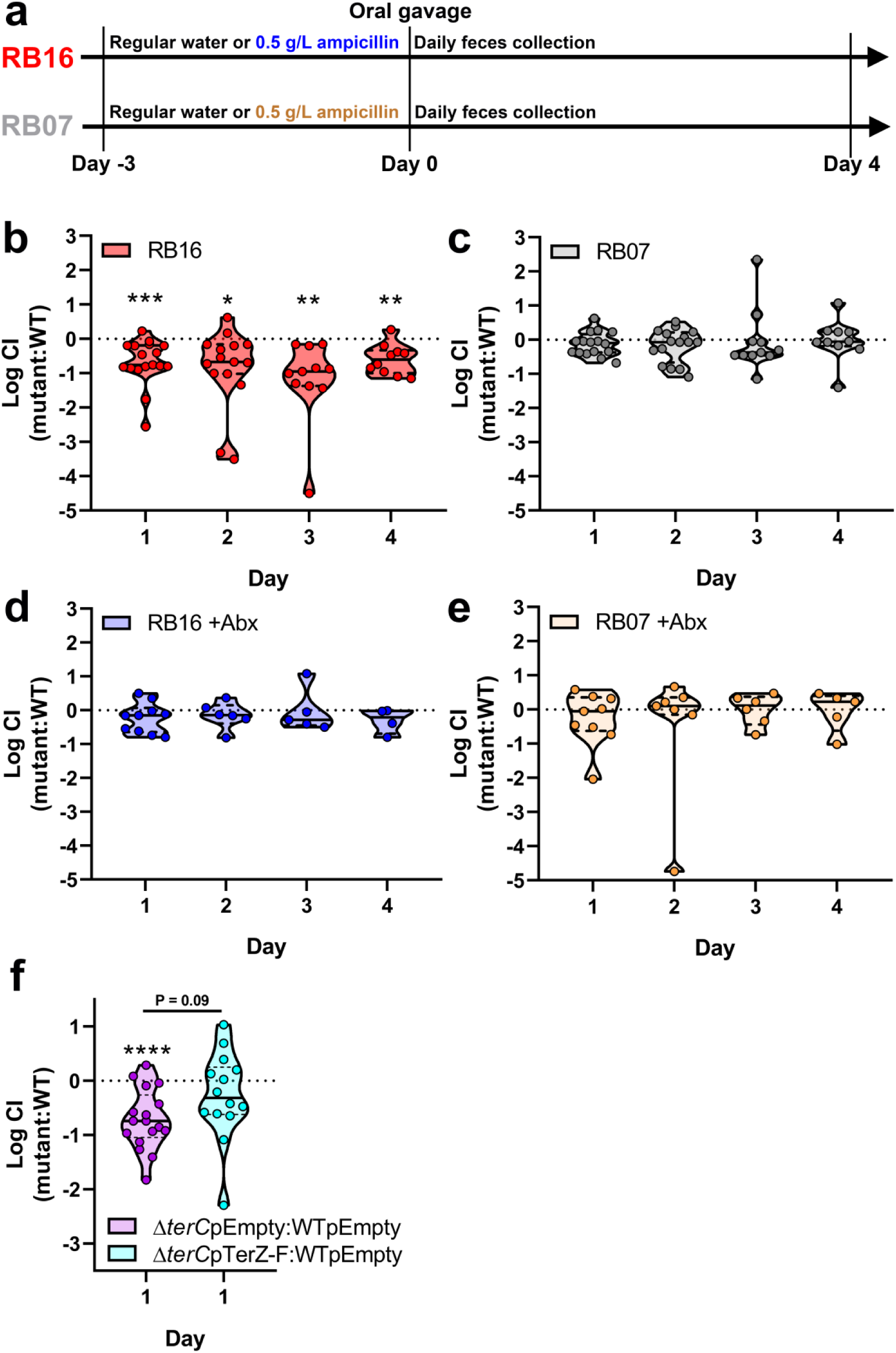
TerC is a fitness factor during gut colonization. **a**, Three days prior to inoculation, male and female C57BL6/J mice sourced from barriers RB16 and RB07 were treated with 0.5 g/L ampicillin or regular drinking water. **b**-**e**, NTUH-K2044 and the isogenic Δ*terC* mutant were mixed 1:1 and approximately 5×10^6^ CFU were orally gavaged into mice (n = 9-18 per group). A fresh fecal pellet was collected daily from each animal, CFUs were enumerated, and log competitive indices (mutant:WT) were calculated (median and IQR displayed, *P < 0.05, **P < 0.005, ***P < 0.0005, one-sample *t* test compared to a hypothetical value of 0). **f**, NTUH-K2044 and the isogenic Δ*terC* mutant containing an empty vector or the pTerZ-F plasmid were mixed 1:1 and approximately 5×10^6^ CFU were orally gavaged into mice sourced from barrier RB16 (n = 14-16). A fresh fecal pellet was collected 24 hours after inoculation, CFUs were enumerated, and log competitive indices (mutant:WT) were calculated (f, median and IQR displayed, ****P < 0.00005, one-sample *t* test compared to a hypothetical value of 0 or Student’s *t* test). Each data point represents an individual animal.

**Fig. 4:**
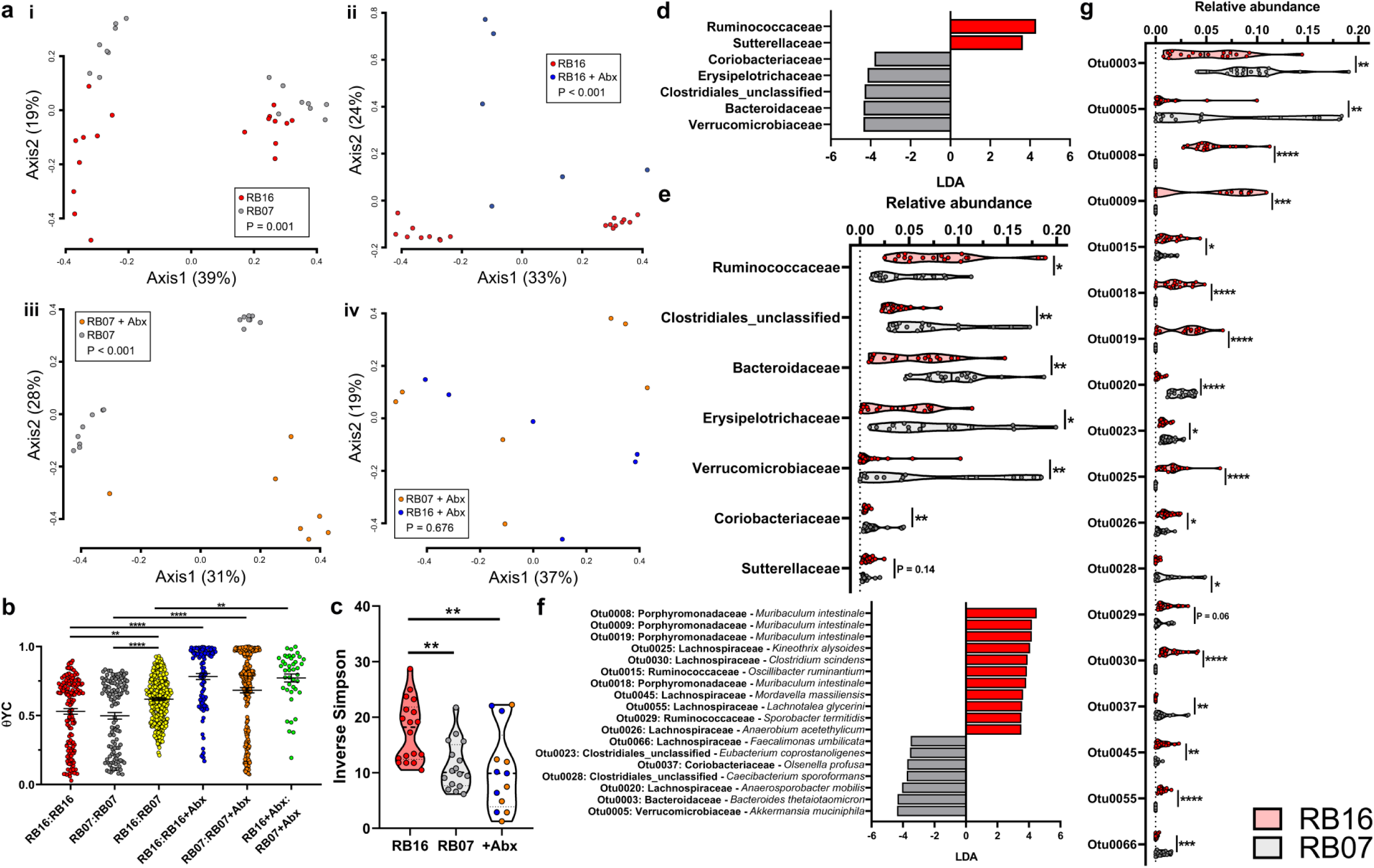
The fecal microbiota in which *terC* is (RB16) and is not (RB07) a fitness factor are distinct. Fecal pellets collected from male and female C57BL6/J mice sourced from barriers RB16 and RB07 (n = 9-20 mice per group) on the day of Kp inoculation were subjected to 16S rRNA gene sequencing. **a**,**b**, Pairwise community dissimilarity values between the fecal microbiota communities of barriers RB16 and RB07 with or without three days treatment with 0.5 g/L ampicillin were visualized by Principal coordinates analysis (**a**, AMOVA) and individually (**b**, **P < 0.005, ****P < 0.00005, one-way ANOVA followed by Tukey’s multiple comparisons post-hoc test). **c**, Diversity of the fecal microbiota was summarized by inverse Simpson index (**P < 0.005, one-way ANOVA followed by Tukey’s multiple comparisons post-hoc test). **d**,**f**, LEfSe was used to determine if specific bacterial families (**d**) or OTUs (**f**) were differentially abundant between the fecal microbiota of RB16 and RB07 (**d**, LDA ≥ 3.5 and P < 0.05 are shown). **e, g**, differential bacterial families (**e**) or OTUs (**g**) relative abundance values were plotted (**e**, *P < 0.05, **P < 0.005, ***P < 0.0005, ****P < 0.00005, Student’s *t* test).

To determine the composition of the gut microbiota of mice from RB16 in which Δ*terC* had reduced fitness, and RB07 in which it did not, we performed sequence analysis of amplicons of the 16S rRNA-encoding gene from fecal DNA collected throughout the course of these experiments (Supplementary Fig. 7, Supplementary Tables 4, 5). To compare the microbiota between all groups (RB16, RB07, RB16+Abx, RB07+Abx) and all time points, θ_YC_ distances^27^ were calculated, and principal coordinates analysis was used to visualize these distances. As expected^28-30^, microbiota differed between male and female mice (Fig. 4a.i, axis 1, females cluster on left of graph). Despite this, on the day of inoculation the fecal microbiota of RB16 and RB07 were significantly dissimilar (Fig. 4a.i, axis 2, AMOVA P = 0.001; Fig. 4b), suggesting that the results observed in Fig. 3b-c were attributable to differences in the microbiota of these mice. In addition, the fecal microbiota of RB16 and RB16+Abx (Fig. 4a.ii, AMOVA P < 0.001), as well as RB07 and RB07+Abx, were significantly dissimilar (Fig. 4a.iii, AMOVA P < 0.001); however, the microbiota of RB16+Abx and RB07+Abx were not significantly dissimilar (Fig. 4a.iv, AMOVA P = 0.676). Assessment of the magnitude of dissimilarity indicated that intergroup dissimilarity was higher than intragroup dissimilarity, and antibiotic treatment resulted in the greatest dissimilarity (Fig. 4b). These findings were consistent across all time points (Supplementary Fig. 8). These data demonstrate an association between *ter*-dependent fitness in the gut and the composition of the gut microbiota, and suggest that an individual or group of gut microbiota constituents in the RB16 background underlies the observed loss of fitness.

There were a number of differences between the microbiota in mice RB16 and RB07 mice. The diversity of the fecal microbiota of RB16 was significantly higher than RB07 at the day of inoculation (Fig. 4c), and this difference was present throughout the course of the experiment (Supplementary Fig. 9). We next sought to determine if a bacterial family or families differentiated the fecal microbiota of RB16 and RB07. To this end, we used linear discriminant analysis (LDA) effect size (LEfSe)^31^ to determine if specific families were differentially abundant between RB16 and RB07. On the day of inoculation, 7 bacterial families were found to be differentially abundant, 2 of which were more abundant in RB16 and 5 of which were more abundant in RB07 (Fig. 4d-e), and of these, only unclassified Clostridiales associated with RB07 remained differential throughout the entire experiment (Supplementary Figs. 10-11). We hypothesized that no family consistently distinguished RB16 from RB07 because variations in individual OTU relative abundances obscured family-level analysis. As such, we used LEfSe analysis to determine differentially abundant operational taxonomic units (OTUs) between these microbiotas. On the day of inoculation, 18 OTUs were found to be differentially abundant, 7 of which were more abundant in RB07, and 11 of which were more abundant in RB16 (Fig. 4f). Intriguingly, 4 of the 11 OTUs associated with barrier RB16 had 16S rRNA sequences most similar to *Muribaculum intestinale*^32-34^ (Fig. 4f; 0008, 0009, 0018, 0019). These 4 OTUs, as well as 0030 which had 16S rRNA sequence most similar to *Clostridium scindens*, remained more abundant in RB16 than RB07 for the entire experiment (Supplementary Figs. 12-13). Notably, the Porphyromonadaceae family, of which *M. intestinale* was considered a member in our dataset, was also enriched in RB16 on days 1-3 post-inoculation (Supplementary Figs. 10-11). The identification of these species as differentially abundant in RB16 is notable, as *M. intestinale* belongs family S24-7 which is suggested to play a role in gut inflammatory homeostasis^22,34-36^, and *C. scindens* has been reported to be a clinically relevant probiotic candidate^37^. The only OTU that was more abundant in mice sourced from barrier RB07 throughout the course of the experiment is 0037, which is most similar to *Olsenella profusa* (Fig. 4f, Supplementary Figs. 12-13). Notably, OTUs enriched in RB16 were mostly absent in RB07 (Fig. 4g) throughout the course of the experiment (Supplementary Figs. 12-13). If these OTUs are tightly associated with the observed *terC* fitness defect, they should also be sensitive to the antibiotic treatment that ameliorates the defect. Indeed, differential families (Supplementary Figs. 14-15) and OTUs (Supplementary Figs. 16-17) were sensitive to antibiotic treatment highlighting their potential interaction with *ter*. Finally, we asked if the introduction of Kp impacted the relative abundance of these families and OTUs. At the family level, the microbiota of barrier RB16 were only minimally modulated (Supplementary Fig. 18a) by Kp inoculation, and the only OTU that differentiated RB16 from RB07 that was modulated was 0045 (*Mordavella massiliensis*, Supplementary Fig. 18b). These data indicate that the microbiota constituents of RB16 that interact with *ter* are highly stable, and suggest that the interaction is either indirect or that it impacts Δ*terC* fitness without modulation of relative abundance.

Taken together, our data demonstrates that the Kp *ter* operon is important for gut fitness in a microbiome-dependent manner. The biological function of *ter* has long been enigmatic, as TeO_3_^−2^ is absent in humans. Conversely, the *ter* locus has been identified in multiple pathogens^10,38-40^, including Kp. This study provides a plausible biological function of *ter* that could directly explain its association with clinical infection. Gut colonization is a critical first step for many pathogens that cause both intestinal and extra-intestinal infection. The *ter* locus is needed for optimal fitness in the presence of certain indigenous gut microbes, suggesting that acquisition of *ter+* plasmids can expand the host range of a pathogen by resisting the competitive pressures of these bacteria. This evasion of colonization resistance by horizontal gene transfer echoes the bacterial arms race seen in response to nutritional immunity and antibiotics. Recently, *ter* was suggested to be a point of recombination for the Kp hypervirulence plasmid and a carbapenemase encoding plasmid^41^, suggesting it can both enhance gut fitness and enable the convergence of two worrying Kp pathotypes. Given that it is expected that by the year 2050 up to 10 million deaths per year will be due to drug-resistant bacteria, resulting in a loss of up to $100 trillion of economic output^42^, studies that identify and characterize bacterial factors associated with pathogenesis are critical. Fortunately, the identification of certain indigenous gut microbes as a significant barrier to gut colonization by potential pathogens reinforces the potential of microbiome editing to eliminate pathogen colonization.

## Methods

### Ethics statement

This study was performed in strict accordance with the recommendations in the *Guide for the Care and Use of Laboratory Animals*^43^. The University of Michigan Institutional Animal Care and Use Committee approved this research (PRO00007474).

### Materials, media, and bacterial strains

All chemicals were purchased from Sigma-Aldrich (St. Louis, MO) unless otherwise indicated. *E. coli* K12 strain MG1655, Kp strain NTUH-K2044^44^, and isogenic mutants were cultured in Luria-Bertani (LB, Becton, Dickinson and Company, Franklin Lakes, NJ) broth at 37°C with shaking, or on LB agar at 27°C (Thermo Fisher Scientific). The isogenic *terC* mutant^1^, and grown in the presence of 40 µg/mL kanamycin. To construct the pTerC and pTerZ-F complementation plasmids, PCR products derived from WT NTUH-K2044 containing the *terC* or *terZABCDEF* open reading frames were inserted into pCR 2.1 using TOPO TA cloning (Life Technologies, Carlsbad, CA) and subsequently ligated into pACYC184 following digestion with Xbal and HindIII. The ligation mixture was transformed into NEB 10-beta Competent *E. coli* (New England Biolabs) by heat shock. *E. coli* transformants were selected at 37°C on LB agar containing 30 µg/ml chloramphenicol, re-cultured, and confirmed by colony PCR. Single transformants were then grown in batch culture for plasmid extraction using the Plasmid Midi Kit (Qiagen, Germantown, MD). MG1655 and NTUH-K2044 competent cells were prepared as previously described^45^, electroporated with the complementation plasmids or corresponding empty vector, and selected at 37°C on LB agar containing 30 (MG1655) or 80 µg/ml (NTUH-K2044) chloramphenicol. Following selection, transformants were re-cultured, and confirmed by colony PCR and by growth in presence of TeO_3_^−2^. All primer sequences can be found in Supplementary Table 6. For all subsequent experiments, complemented and control strains were grown in the presence of the appropriate antibiotic.

### Plasmid sequencing and analysis

To characterize *ter*-encoding Kp strains, the *wzi* gene sequence was extracted from each isolate and Wzi type was assigned using the Bacterial Isolate Genome Sequence Database (BIGSdb)^17,46^. To characterized the *ter*-encoding plasmids from our previous study^1^, genomic DNA was extracted from pure Kp cultures using the DNeasy PowerSoil Pro Kit (Qiagen, Hilden, Germany). Long-read genomic sequencing was performed using GridION X5 (Oxford, England) sequencing instrument. Each Nanopore sequencing library was prepared using 1µg of DNA with the 1D ligation kit (SQK-LSK108, Oxford Nanopore Technologies) and sequenced using R9.4.1 flowcells (FLO-MIN106). MinKNOW software was used to collect sequencing data. Nanopore reads were called using Albacore v2.2.3, and assembled using Canu v1.7^47^. Assemblies were corrected for ten rounds with Illumina reads using Pilon v1.22^48^ in conjunction with the bowtie2 v2.3.3.1 aligner^49^. *ter*-encoding reference strains and plasmids from the NCBI database were identified using BLAST^50^, wherein the entire *ter* locus (Supplementary Table 3) was used as the query, and *Klebsiella pneumoniae* (taxid:573) as the subject (extraction date 03/27/2019). *ter* was not identified on any Kp chromosomes.

Assembled plasmid sequences were circularized and annotated the plasmids using Dfast^51^ prokaryotic annotation pipeline. Pairwise alignments were performed using BLAST^50^. To assess the presence of hvKp and antibiotic resistance genes, reference sequences (Supplementary Table 3) were extracted, and BLAST^50^ was used to align these reference sequences to *ter* encoding plasmid sequences. To study the genes conserved around the *ter* locus, gene level multiple sequence alignment (MSA) of the genes within 10 kbp upstream of the putative biosynthetic locus and downstream of the *ter* operon in all the plasmids. These loci were visualized by coding annotated genes using their 4-character gene names and unannotated hypothetical proteins using their gene cluster identifier as determined by CD-HIT software^52^ for the MSA. The MSA was performed using MAFFT^53^ in the L-INS-i mode and visualized using MSAviewer^54^ to understand conserved genes around the *ter* locus. To predict protein structure and function of genes within and adjacent to the *ter* locus, *ter* encoding plasmids were annotated using the PATRIC RAST tool kit^55,56^. Unique annotation frequencies were calculated and then unique predicted amino acid sequences were assessed using I-TASSER^12-14^. Supplementary Table 1 indicates reference amino acid sequences used for protein structure and function prediction. Only the highest scoring Predicted Biological Process, Predicted Molecular Function, and Predicted Pubchem Ligand Binding Site are reported. Finally, complete plasmid MSA was performed and visualized using Mauve (MegAlign Pro, DNASTAR Inc., Madison, WI).

### Murine models of infection

Six- to 12-week-old C57BL/6J male and female mice from barriers RB07 and RB16 (Jackson Laboratory, Jackson, ME) were used for all murine models of infection. Gender was evenly distributed in all groups. For bacteremia studies, WT NTUH-K2044 and NTUH-K2044Δ*terC* were cultured overnight in LB, then bacteria were pelleted, resuspended, mixed 1:1, diluted in sterile PBS to the appropriate dose, and mice were inoculated intraperitoneally with approximately 5×10^5^ CFU. After 24 hours, mice were euthanized by CO_2_ asphyxiation and blood, spleen, liver, and brain were collected. Solid organs were weighed and homogenized in sterile PBS, and whole blood and solid organ homogenates were plated on selective media. For oral inoculation studies, mice from both barriers were given regular drinking water, or water containing 0.5 g/L ampicillin 3 days prior to inoculation and throughout the experiment. Kp strains were cultured overnight in LB in the presence of antibiotics when appropriate, then bacteria were pelleted, resuspended, mixed 1:1, diluted in sterile PBS to the appropriate dose, and mice were orally inoculated with approximately 5×10^6^ CFU. For four days post-inoculation, a fresh fecal pellet was collected from each mouse, weighed, and homogenized in sterile PBS, and homogenates were dilution plated on both LB agar containing 10 µg/ml ampicillin or 40 µg/ml kanamycin to determine Kp load. When complemented or empty vector control strains were used, plasmid maintenance was monitored by plating both on LB agar containing 10 µg/ml ampicillin or 40 µg/ml kanamycin to determine total Kp load, and on LB agar containing 10 µg/ml ampicillin and 80 µg/ml chloramphenicol or 40 µg/ml kanamycin and 80 µg/ml chloramphenicol to determine plasmid maintaining Kp load. In all models, mice were monitored daily for signs of distress (hunched posture, ruffled fur, decreased mobility, and dehydration) and euthanized at predetermined timepoints, or if signs of significant distress were displayed. No blinding was performed between experimental groups.

### 16S rRNA gene sequencing and analysis

Fecal DNA was isolated from using the MagAttract PowerMicrobiome DNA/RNA Kit (Qiagen) and an epMotion 5075 liquid handling system. The V4 region of the 16S rRNA gene was amplified and sequenced as previously described^57^. Standard PCRs used 1, 2 or 7 μL of undiluted DNA and touchdown PCR used 7 μL of undiluted DNA. 16S rRNA gene sequence data was processed and analyzed using the software package mothur (v.1.40.2)^58,59^. Sequences were binned into OTUs based on 97% sequence similarity using the OptiClust method^60^ following sequence processing and alignment to the SILVA reference alignment (release 128)^61^. θ_YC_ distances^27^ were calculated between communities, and AMOVA^62^ was used to determine statistically significant differences between experimental groups^27^. Principal coordinates analysis was used to visualize the θ_YC_ distances between samples. Taxonomic composition of the bacterial communities was assessed by classifying sequences within mothur using a modified version of the Ribosomal Database Project training set (version 16)^63,64^, and diversity metrics, including inverse Simpson, were calculated. Finally, linear discriminant analysis effect size was used to determine if specific families and OTUs were differentially abundant in different groups^31^. Putative genus and species assignment was performed by comparing the representative 16S rRNA sequences from OTUs to the NCBI 16S ribosomal RNA sequence database. These assignments were confirmed using the Ribosomal Database Project (RDP) database, with the exception of OTUs assigned to *Muribaculum intestinale* based on NCBI, but to Porphyromonadaceae by RDP^63^. All other assignments were in agreement.

## Data availability

Illumina reads are available on the Sequence Read Archive (SRA) in BioProject PRJNA464397 (associated figures and tables: Fig. 1, Supplementary Figs. 1 & 2, and Supplementary Table 2). Any other datasets generated during and/or analyzed in this study are available from the corresponding author on reasonable request.

## Statistical analysis

For *in vitro* studies, two-tailed Student’s *t*-tests or ANOVA followed by Tukey’s multiple comparisons post-hoc test was used to determine significant differences between groups. All *in vitro* experimental replicates represent biological replicates. All animal experiments were repeated at least twice with independent bacterial cultures. Competitive indices ((CFU mutant output/CFU WT output)/(CFU mutant input/CFU WT input))^65^ were log transformed and a one-sample *t-*test was used to determine significant differences from a hypothetical value of 0 and paired ratio *t*-test, or two-tailed Student’s *t-*test as indicated in figure legends was used to determine significant differences between groups. All tests were two-sided, and a P value of less than 0.05 was considered statistically significant for the above experiments, and analysis was performed using Prism 6 (GraphPad Software, La Jolla, CA).

## Acknowledgements

We would like to acknowledge the work performed by The University of Michigan Microbial Systems Molecular Biology Laboratory and The University of Michigan Animal Care and Use staff in support of these studies. Additionally, we would like to thank Harry L.T. Mobley and the members of the Mobley laboratory for their insightful discussion of this work. This work was supported by funding from National Institution of Health (https://www.nih.gov/) grants AI125307 to M.A.B and AI130608 to P.J.S. J.V. was supported by the Molecular Mechanisms of Microbial Pathogenesis training grant (NIH T32 AI007528) and the Postdoctoral Translational Scholar Program (NIH UL1TR002240). The funders had no role in study design, data collection and analysis, decision to publish, or preparation of the manuscript. J.V., C.B., S. R., W.T., M.C.S., T.J.S., V.Y., and M.A.B. designed the experiments and J.V., C.B., S.R., R.H., S.M., Y.B., N.S., and Y.F. performed the experiments and analyzed the data. J.V. and M.A.B. wrote and edited the manuscript.

