## Supplementary Materials for "A plasmid locus associated with *Klebsiella* clinical infections encodes a microbiome-dependent gut fitness factor"

**Supplementary Information**

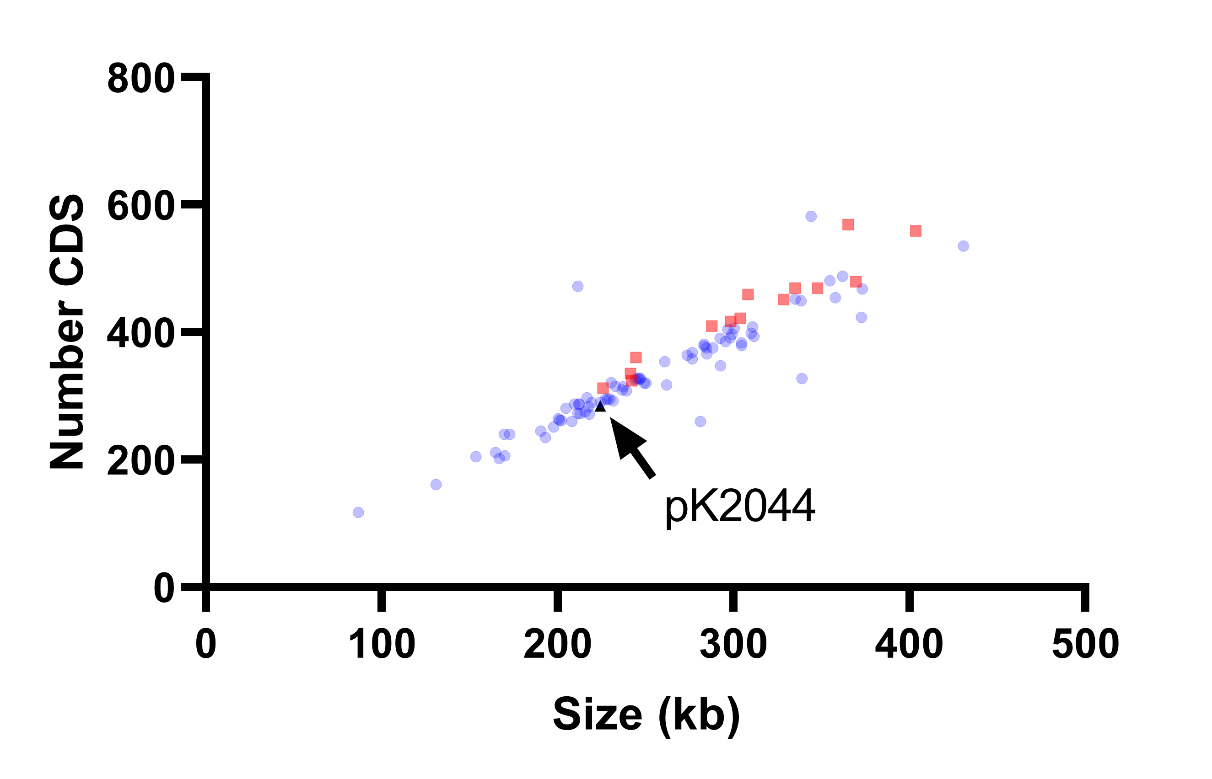

**Supplementary Fig. 1: Size of *ter*-encoding plasmids.**

The size and predicted number of coding sequences (CDS) was determined for plasmids encoding the *ter* operon from Martin *et al.* mSystems, 2018 (red) or reference strains from the NCBI database (blue). The pK2044 hvKp plasmid is shown in black.

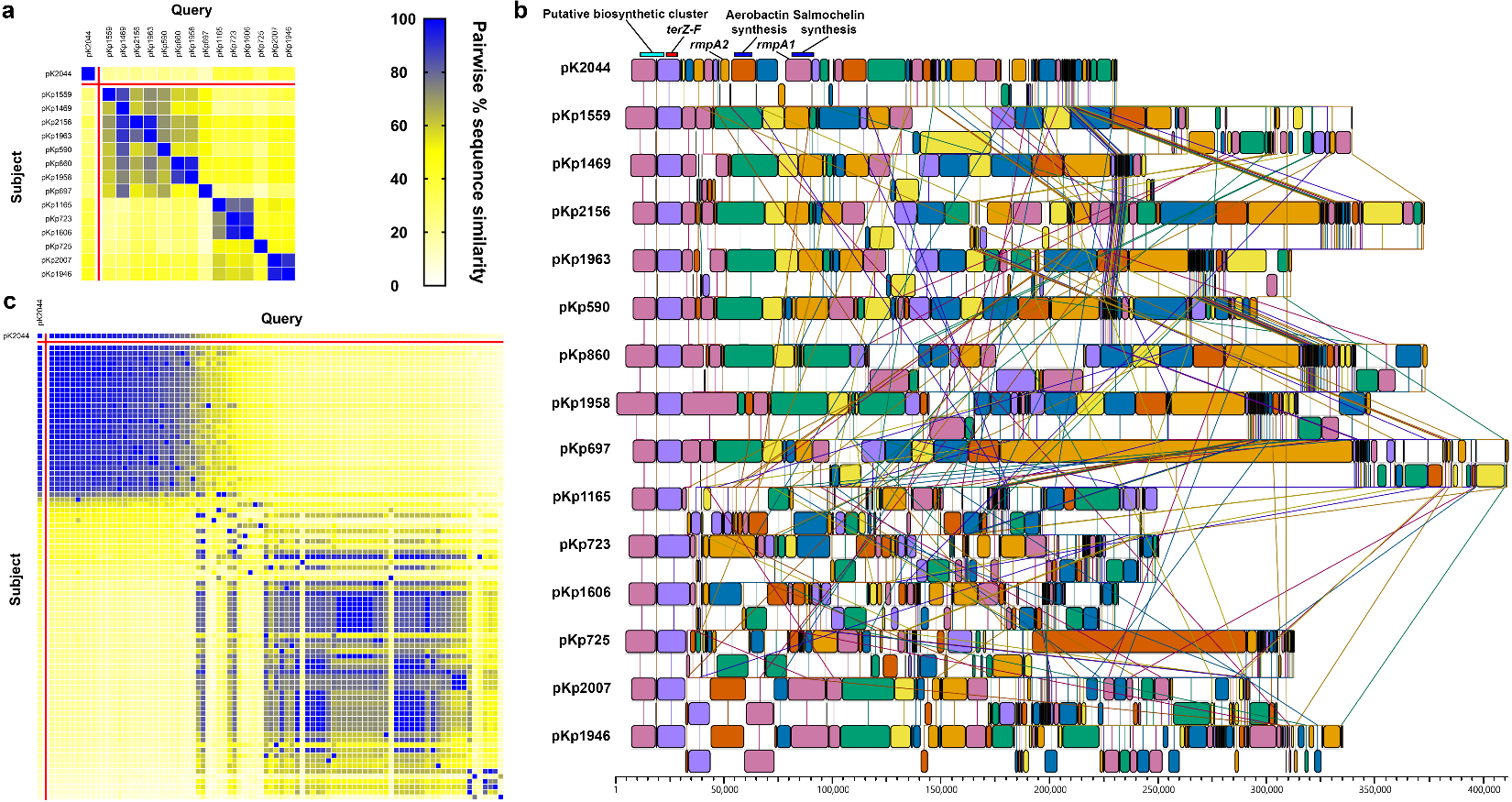

**Supplementary Fig. 2: *ter*-encoding plasmids display variable sequence similarity, gene arrangement and gene content.**

**a**,**b**, Pairwise sequence similarities were determined for plasmids from Martin *et al.* mSystems, 2018 (**a**), and visualized using Mauve (**b**). **c**, Pairwise sequence similarities were also determined for Kp reference plasmids from the NCBI database. **a,c,** Each row and column represents one plasmid. The Kp reference plasmid heat map is organized by pairwise similarity to the pK2044 hvKp plasmid.

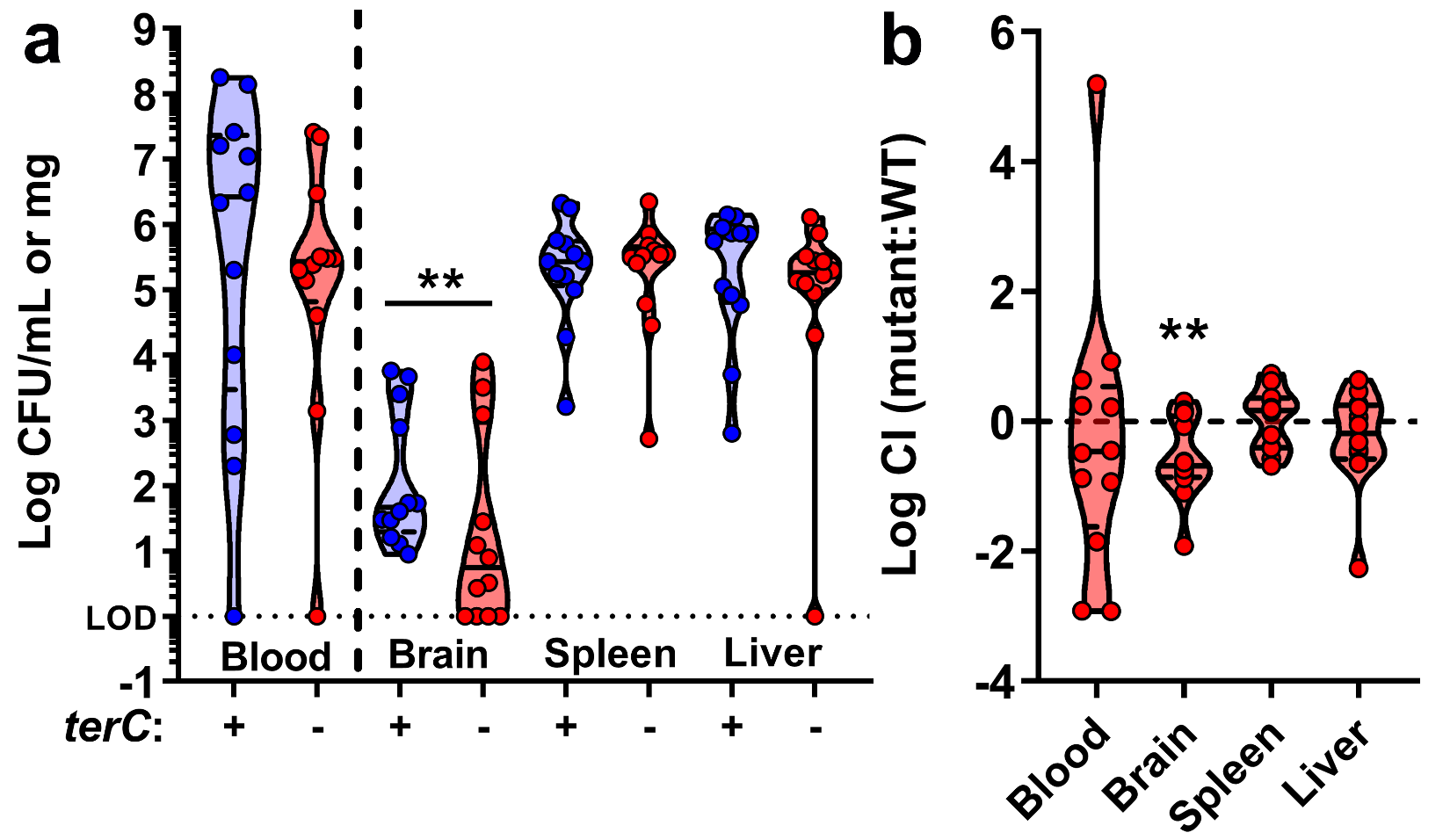

**Supplementary Fig. 3: *terC* is dispensable during bacteremia.**

**a**, **b**, NTUH-K2044 and the isogenic Δ*terC* mutant were mixed 1:1 and approximately 5x10^5^ CFU were inoculated into male and female C57BL6/J mice via peritoneal infection (n = 12). 24 hours post-inoculation, mice were euthanized, tissue CFUs were enumerated (**a**, mean displayed, *P < 0.05, unpaired t test), and log competitive indices (mutant:WT) were calculated (**b**, mean displayed, **P < 0.005, one-sample t test compared to a hypothetical value of 0). Each data point represents an individual animal.

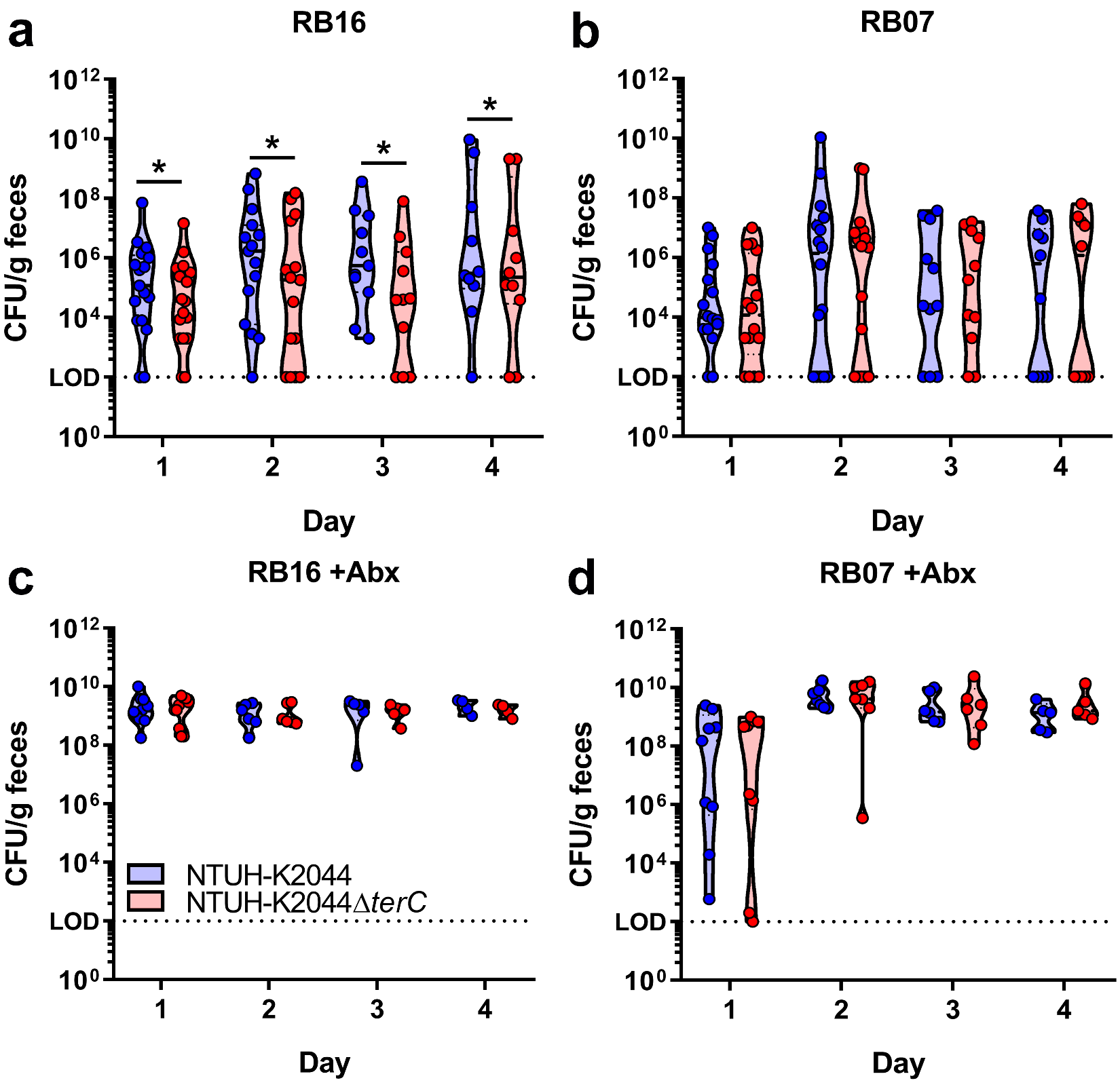

**Supplementary Fig. 4: Gut Kp load during competitive gut colonization.**

**a**-**d**, Three days prior to inoculation, male and female C57BL6/J mice sourced from barriers RB16 and RB07 were treated with 0.5 g/L ampicillin or regular drinking water (Fig. 3a). NTUH-K2044 and the isogenic Δ*terC* mutant were mixed 1:1 and approximately 5x10^6^ CFU were orally gavaged into mice (n = 9-18 per group). A fresh fecal pellet was collected daily from each animal and CFUs were enumerated (median and IQR displayed, *P < 0.05, ratio paired t test).

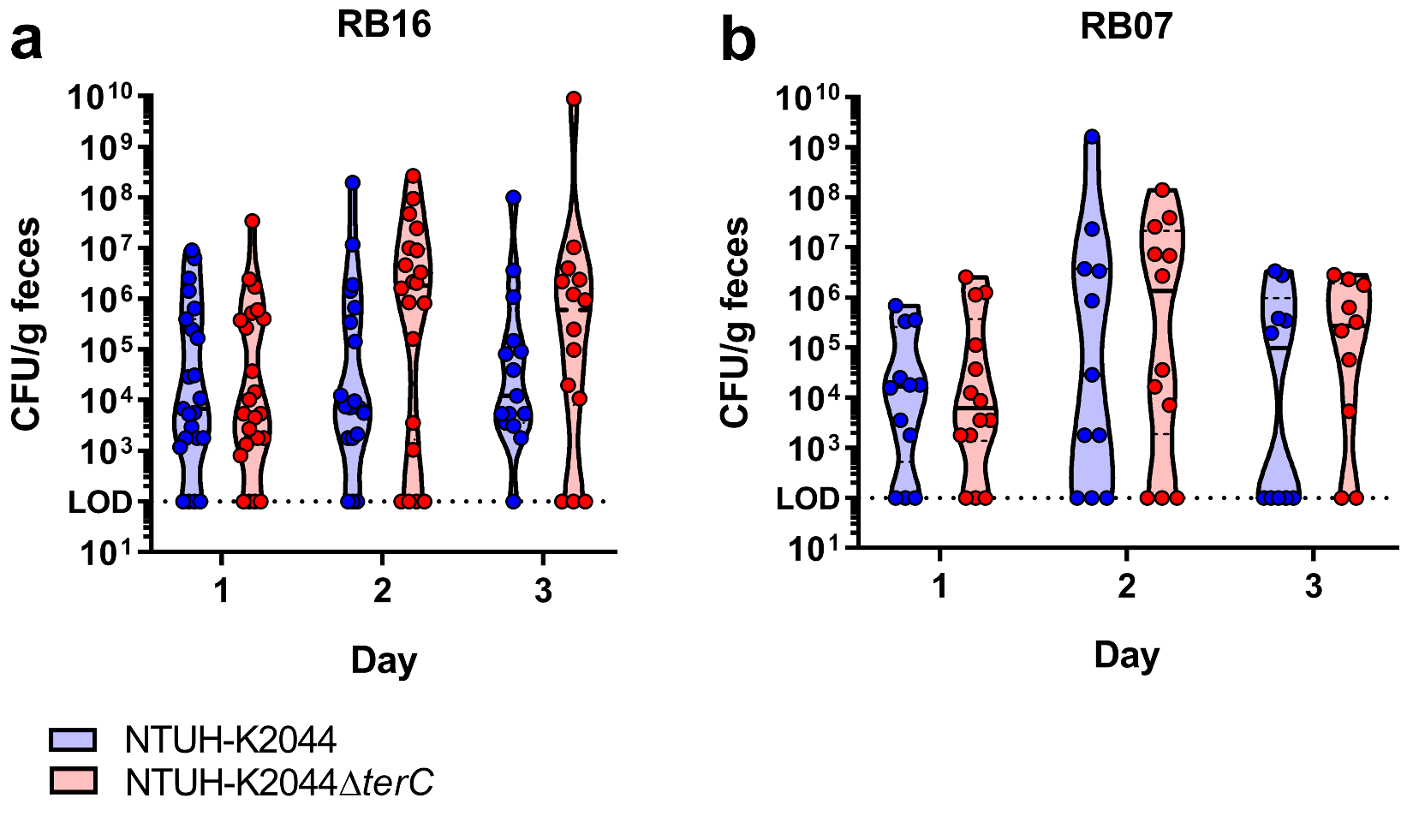

**Supplementary Fig. 5: *terC* is not a colonization factor during mono-strain gut colonization.**

**a**,**b**, Male and female C57BL6/J mice sourced from barriers RB16 (**a**) and RB07 (**b**) were orally gavaged with approximately 5x10^6^ CFU of NTUH-K2044 or the isogenic Δ*terC* mutant (n = 14-24 per group). A fresh fecal pellet was collected daily from each animal and CFUs were enumerated (median and IQR displayed). Each data point represents an individual animal.

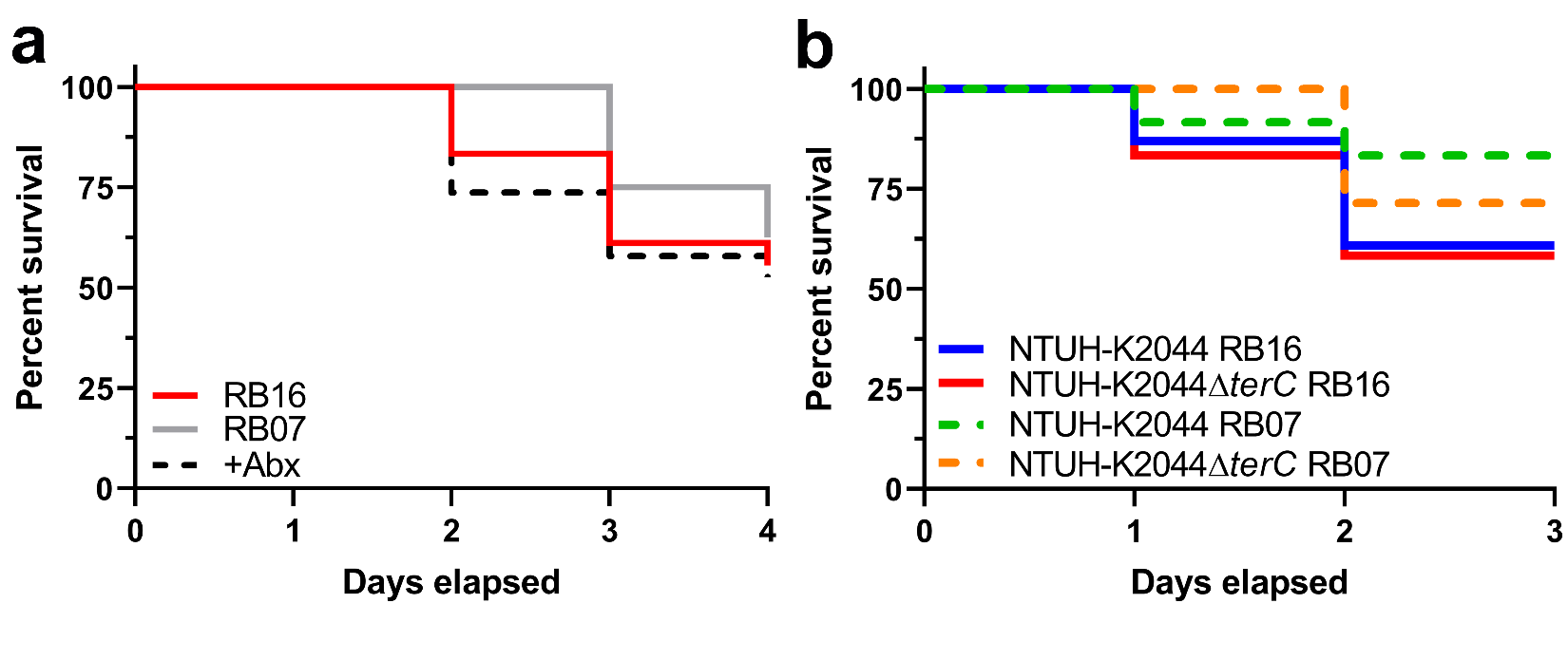

**Supplementary Fig. 6: Mouse survival during gut colonization**

**a**,**b**, Survival of mice from male and female C57BL6/J barriers RB16 and RB07 (16-20 per group) following oral gavage with approximately 5x10^6^ CFU of a 1:1 mix of NTUH-K2044 and the isogenic Δ*terC* mutant (**a**). Survival of mice from male and female C57BL6/J barriers RB07 and RB16 (n = 14-24 per group) following oral gavage with approximately 5x10^6^ CFU of NTUH-K2044 or the isogenic Δ*terC* mutant (**b**). Data were analyzed by Mantel-Cox test between each treatment group in **a** and between each treatment group and between WT and *ΔterC* treated groups in **b**.

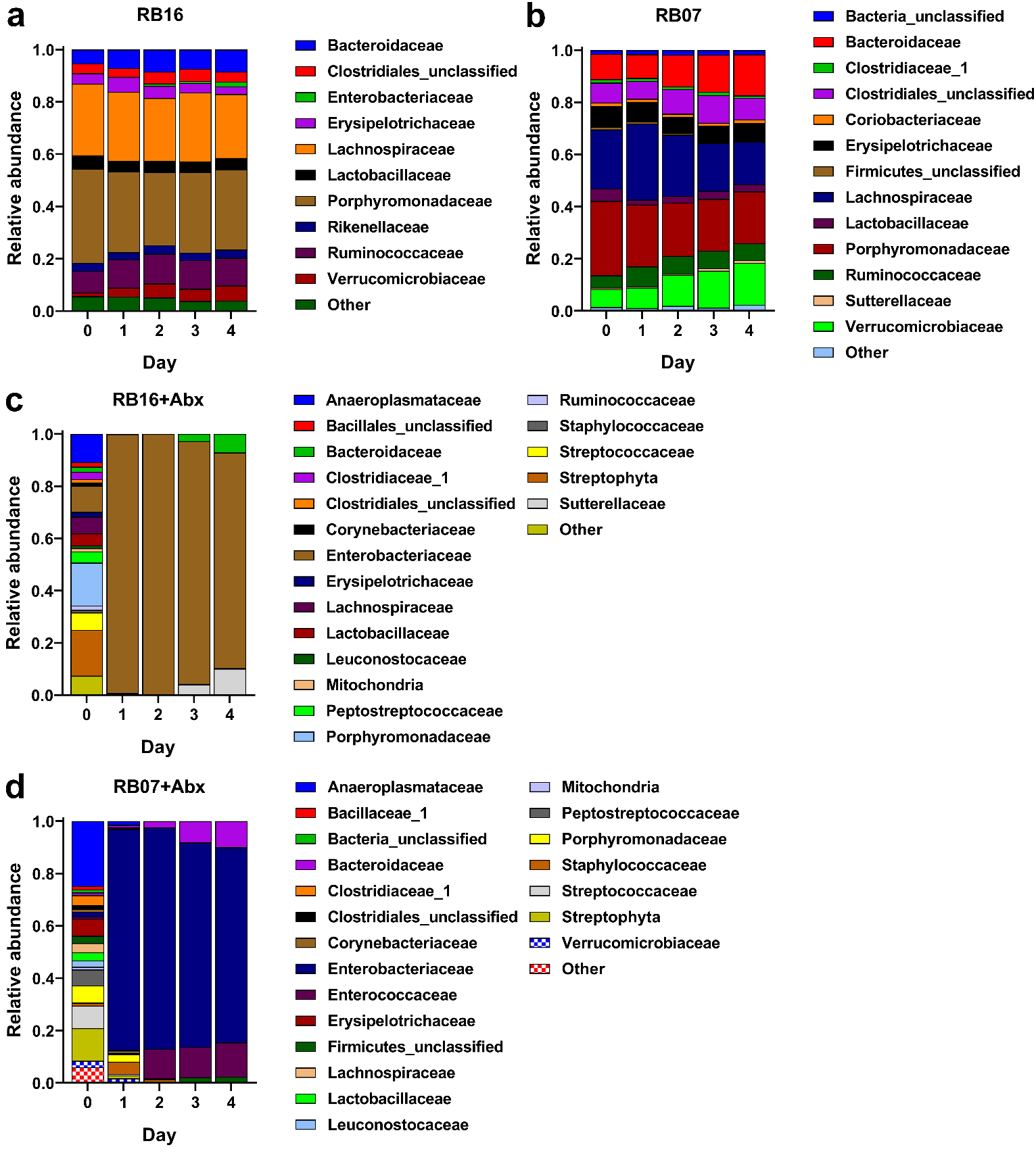

**Supplementary Fig. 7: Community composition of mouse microbiota over time.**

**a**-**d**, Fecal pellets collected daily from male and female C57BL6/J mice sourced from barriers RB16 and RB07 with or without three days treatment with 0.5 g/L ampicillin (n = 9-20 mice per group) following Kp inoculation were subjected to 16S rRNA gene sequencing. Average relative abundance values for bacterial families where relative abundance values are greater than 0.01 are displayed.

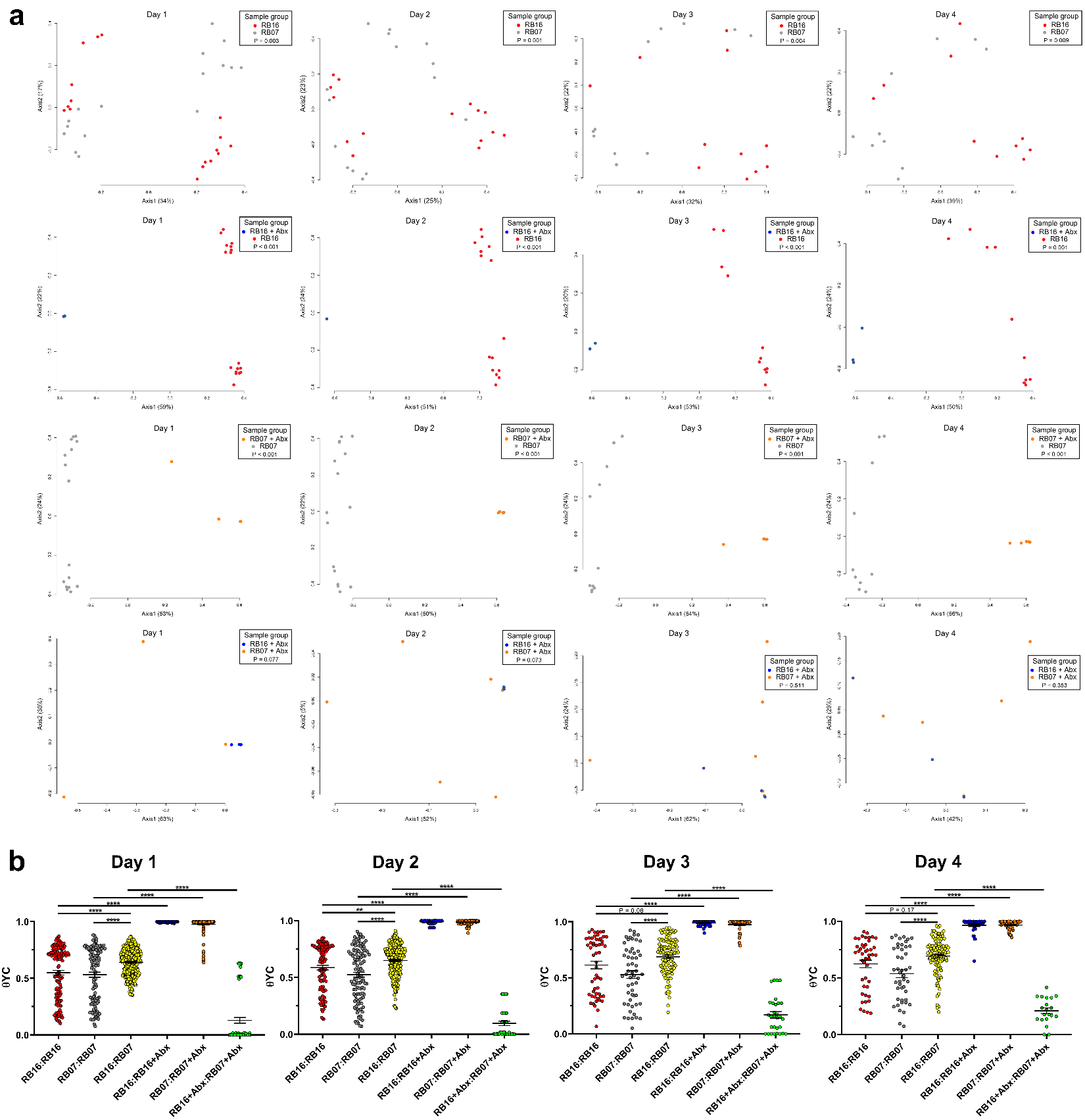
**Supplementary Fig. 8: Differences in community composition of mouse microbiota remain stable over time.**

**a**,**b**, Fecal pellets collected daily from male and female C57BL6/J mice sourced from barriers RB16 and RB07 with or without three days treatment with 0.5 g/L ampicillin (n = 9-20 mice per group) following Kp inoculation were subjected to 16S rRNA gene sequencing. Pairwise community dissimilarity values between the fecal microbiota communities were visualized by Principal coordinates analysis (**a**, AMOVA) and individually (**b**, **P < 0.005, ****P < 0.00005, one-way ANOVA followed by Tukey’s multiple comparisons post-hoc test). Each data point represents an individual animal (**a**) or an individual comparison (**b**).

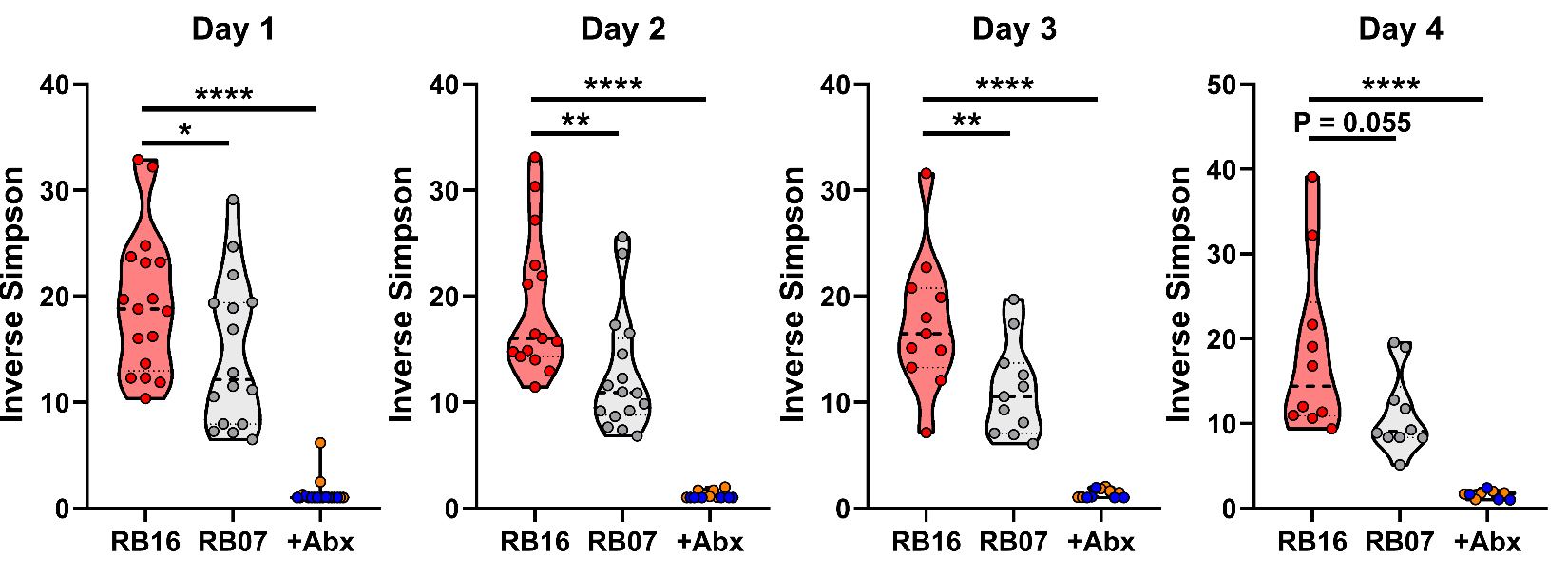
 **Supplementary Fig. 9: Differences in community diversity of mouse microbiota remain stable over time.**

Fecal pellets collected daily from male and female C57BL6/J mice sourced from barriers RB16 and RB07 with or without three days treatment with 0.5 g/L ampicillin (n = 9-20 mice per group) following Kp inoculation were subjected to 16S rRNA gene sequencing. Diversity of the fecal microbiota was summarized by inverse Simpson index (*P < 0.05, **P < 0.005, ****P < 0.00005, one-way ANOVA followed by Tukey’s multiple comparisons post-hoc test). Each data point represents an individual animal. RB16 +Abx is displayed in blue, and RB07 +Abx is displayed in orange.

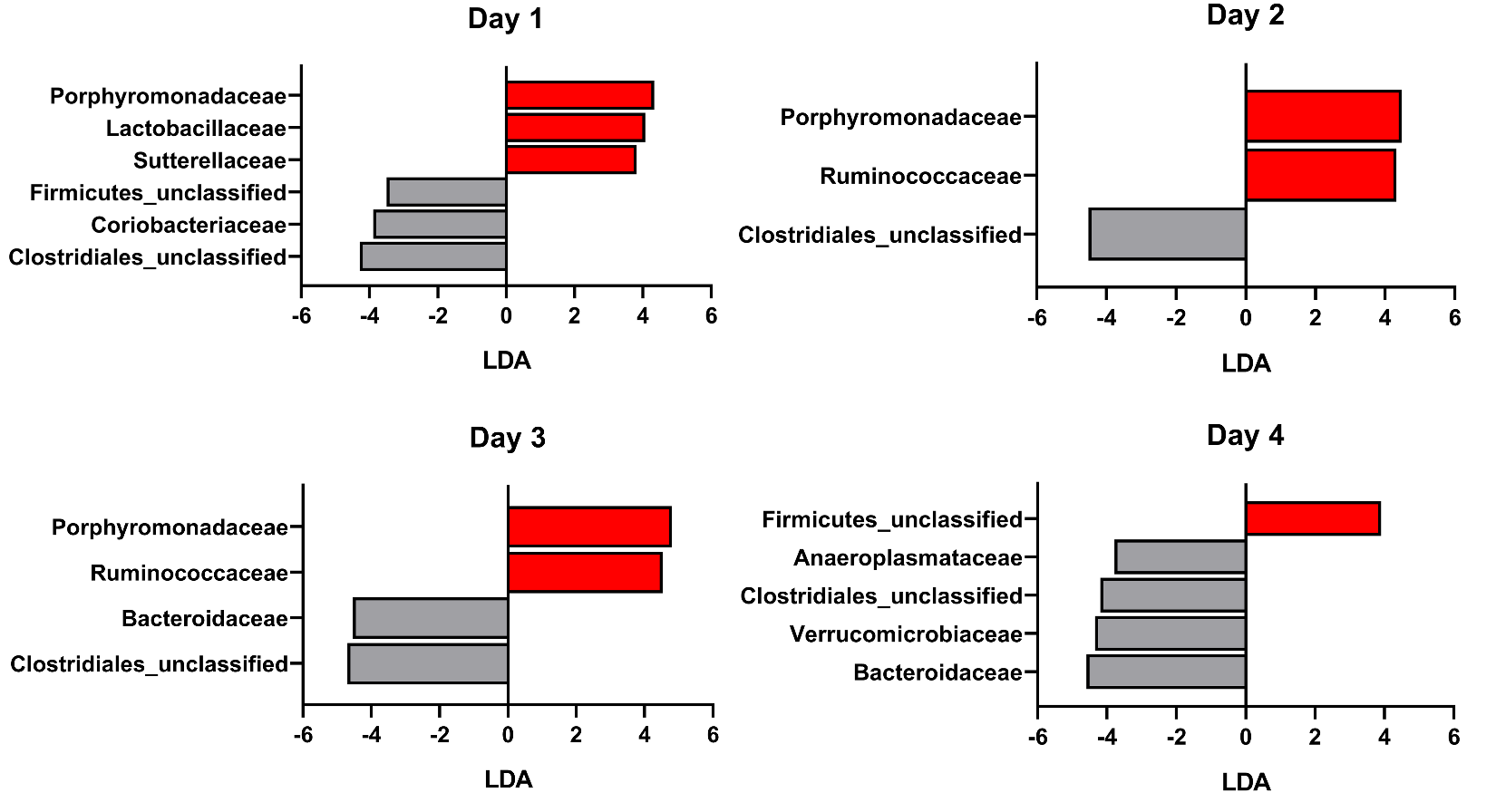
**Supplementary Fig. 10: Differences in bacterial families that differentiate the microbiota of RB16 and RB07 over time.**

Fecal pellets collected daily from male and female C57BL6/J mice sourced from barriers RB16 and RB07 (n = 16-18 mice per group) following Kp inoculation were subjected to 16S rRNA gene sequencing. LEfSe was used to determine if specific bacterial families were differentially abundant between the fecal microbiota of RB16 and RB07 (Families with LDA ≥ 3.5 and P < 0.05 are shown).

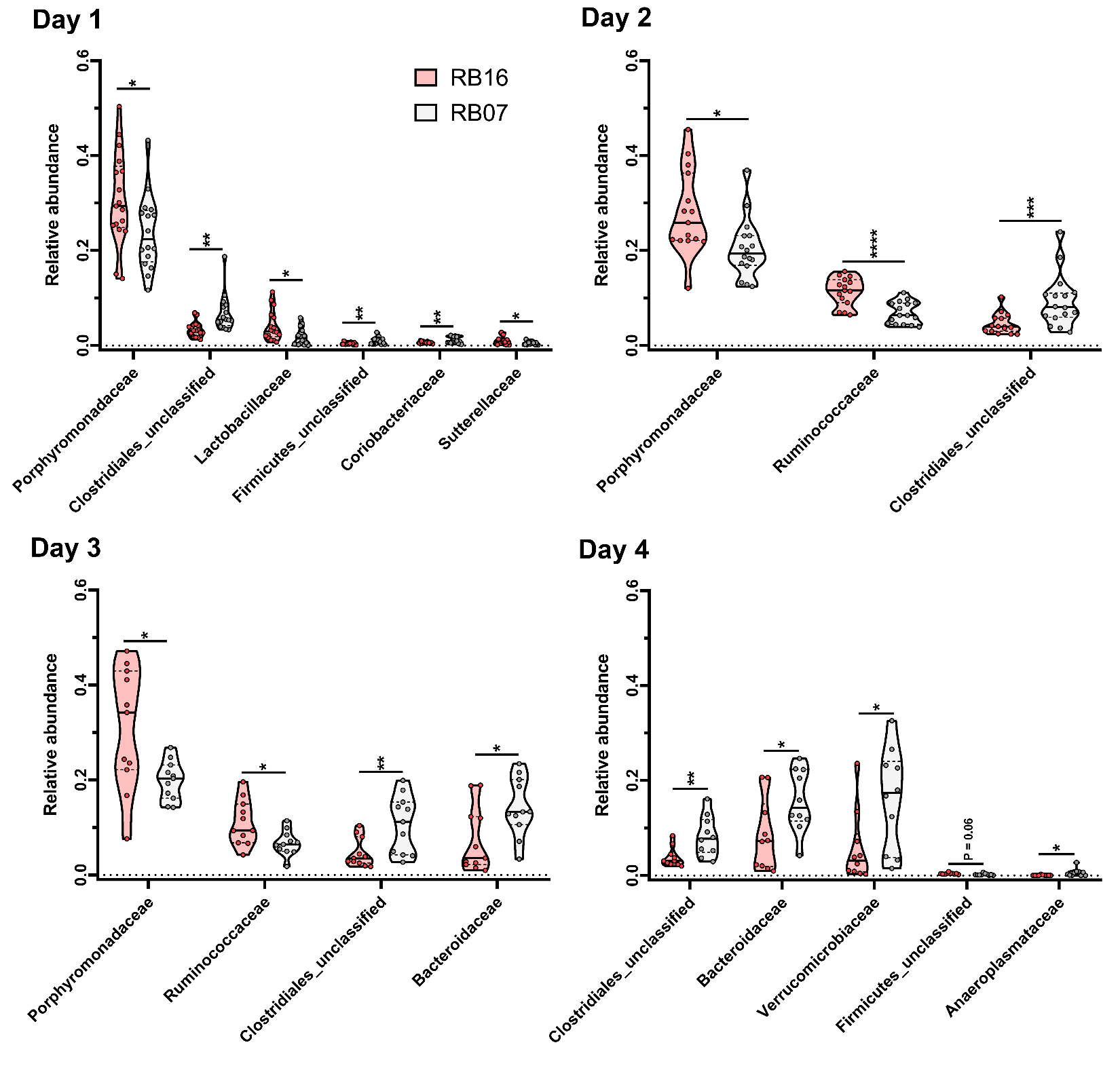
**Supplementary Fig. 11: Differences in relative abundance of bacterial families that differentiate the microbiota of RB16 and RB07 over time.**

Fecal pellets collected daily from male and female C57BL6/J mice sourced from barriers RB16 and RB07 (n = 16-18 mice per group) following Kp inoculation were subjected to 16S rRNA gene sequencing. Relative abundance of specific bacterial families that were differentially abundant between the fecal microbiota of RB16 and RB07 by LEfSe are displayed (*P < 0.05, **P < 0.005, ***P < 0.0005, ****P < 0.00005, Student’s *t* test).

**Supplementary Fig. 12: Differences in OTUs that differentiate the microbiota of RB16 and RB07 remain stable over time.**

Fecal pellets collected daily from male and female C57BL6/J mice sourced from barriers RB16 and RB07 (n = 16-18 mice per group) following Kp inoculation were subjected to 16S rRNA gene sequencing. LEfSe was used to determine if specific OTUs were differentially abundant between the fecal microbiota of RB16 and RB07 (OTUs with LDA ≥ 3.5 and P < 0.05 are shown).

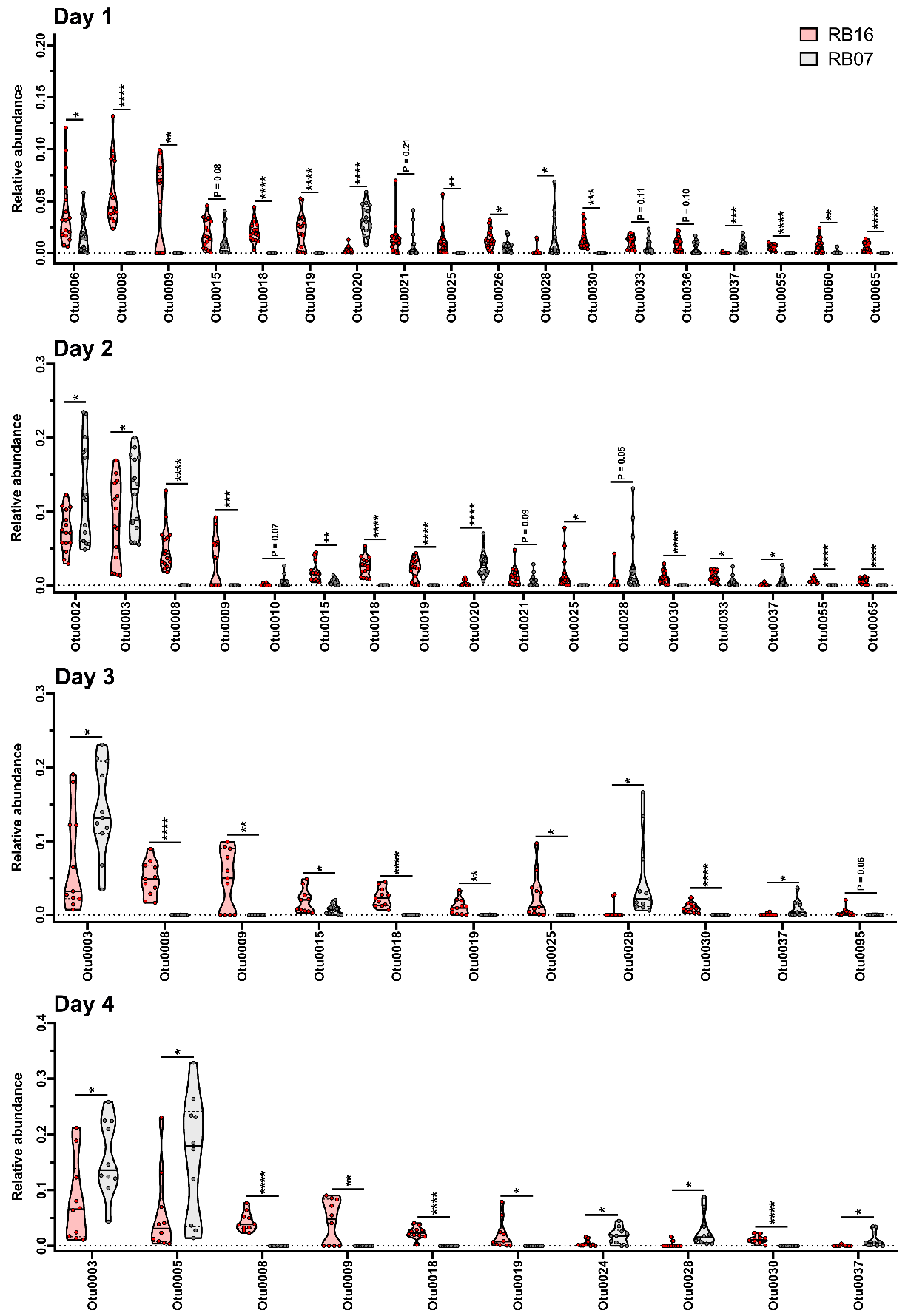

**Supplementary Fig. 13: Differences in relative abundance of OTUs that differentiate the microbiota of RB16 and RB07 remain stable over time.**

Fecal pellets collected daily from male and female C57BL6/J mice sourced from barriers RB16 and RB07 (n = 16-18 mice per group) following Kp inoculation were subjected to 16S rRNA gene sequencing. Relative abundance of specific OTUs that were differentially abundant between the fecal microbiota of RB16 and RB07 by LEfSe are displayed (*P < 0.05, **P < 0.005, ***P < 0.0005, ****P < 0.00005, Student’s *t* test).

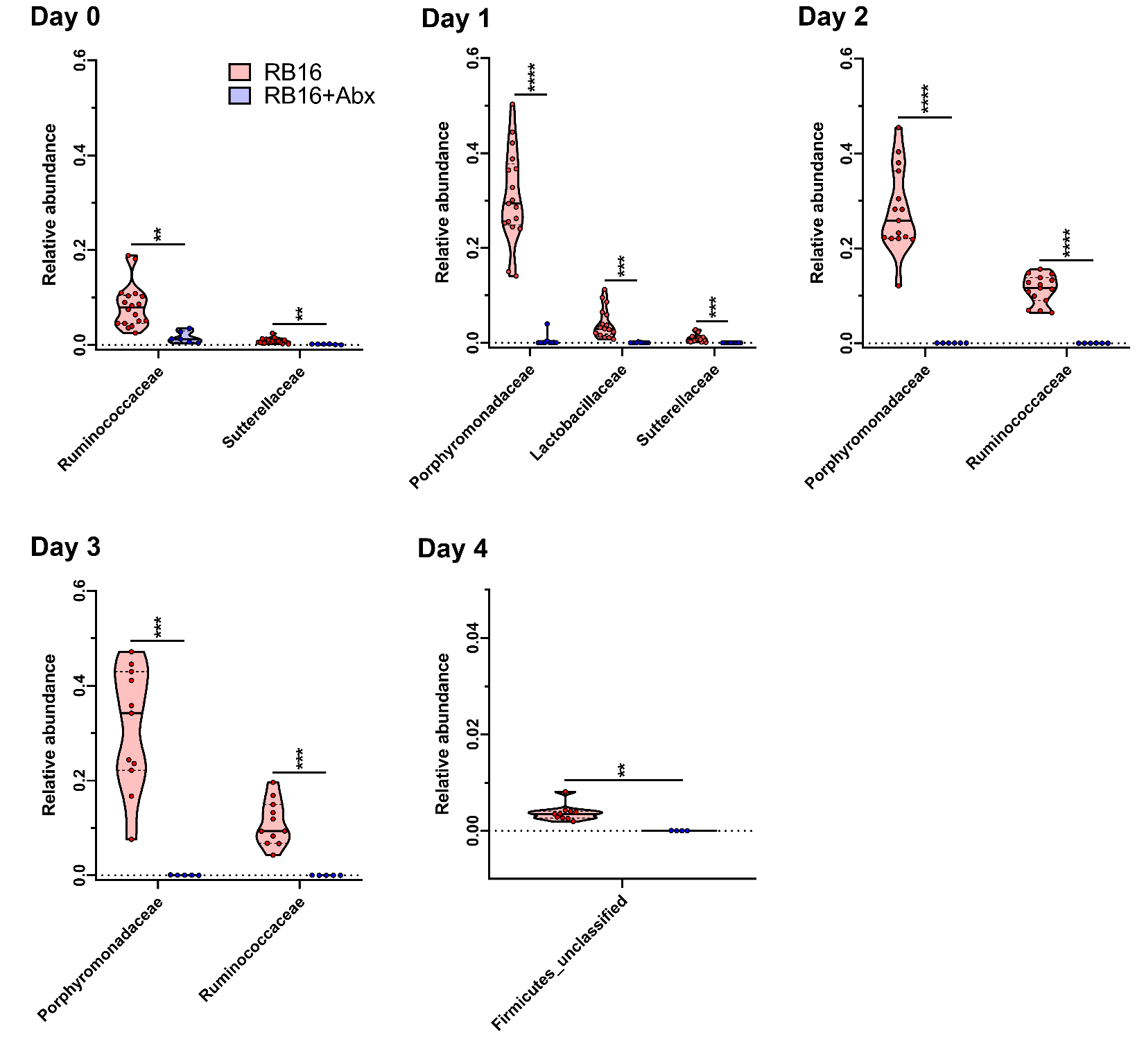
**Supplementary Fig. 14: Bacterial families that differentiate the microbiota of RB16 from RB07 are sensitive to antibiotic treatment.**

Fecal pellets collected daily from male and female C57BL6/J mice sourced from barriers RB16 and RB16+Abx (n = 10-18 mice per group) following Kp inoculation were subjected to 16S rRNA gene sequencing. Relative abundance of specific bacterial families that were differentially abundant between the fecal microbiota of RB16 and RB07 by LEfSe are displayed (**P < 0.005, ***P < 0.0005, ****P < 0.00005, Student’s *t* test).

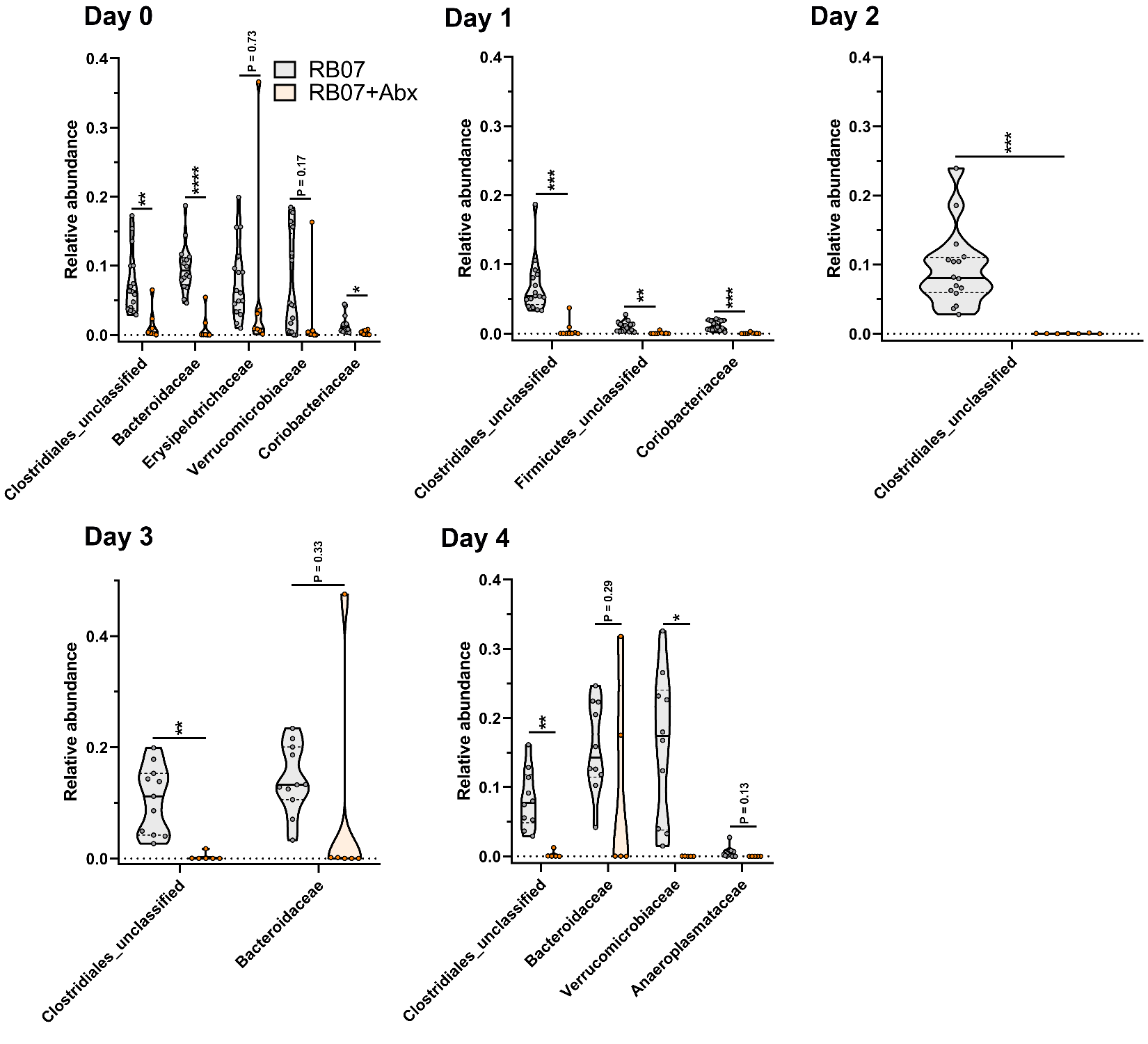
**Supplementary Fig. 15: Bacterial families that differentiate the microbiota of RB07 from RB16 are sensitive to antibiotic treatment.**

Fecal pellets collected daily from male and female C57BL6/J mice sourced from barriers RB07 and RB07+Abx (n = 9-16 mice per group) following Kp inoculation were subjected to 16S rRNA gene sequencing. Relative abundance of specific bacterial families that were differentially abundant between the fecal microbiota of RB16 and RB07 by LEfSe are displayed (*P < 0.05, **P < 0.005, ***P < 0.0005, ****P < 0.00005, Student’s *t* test).

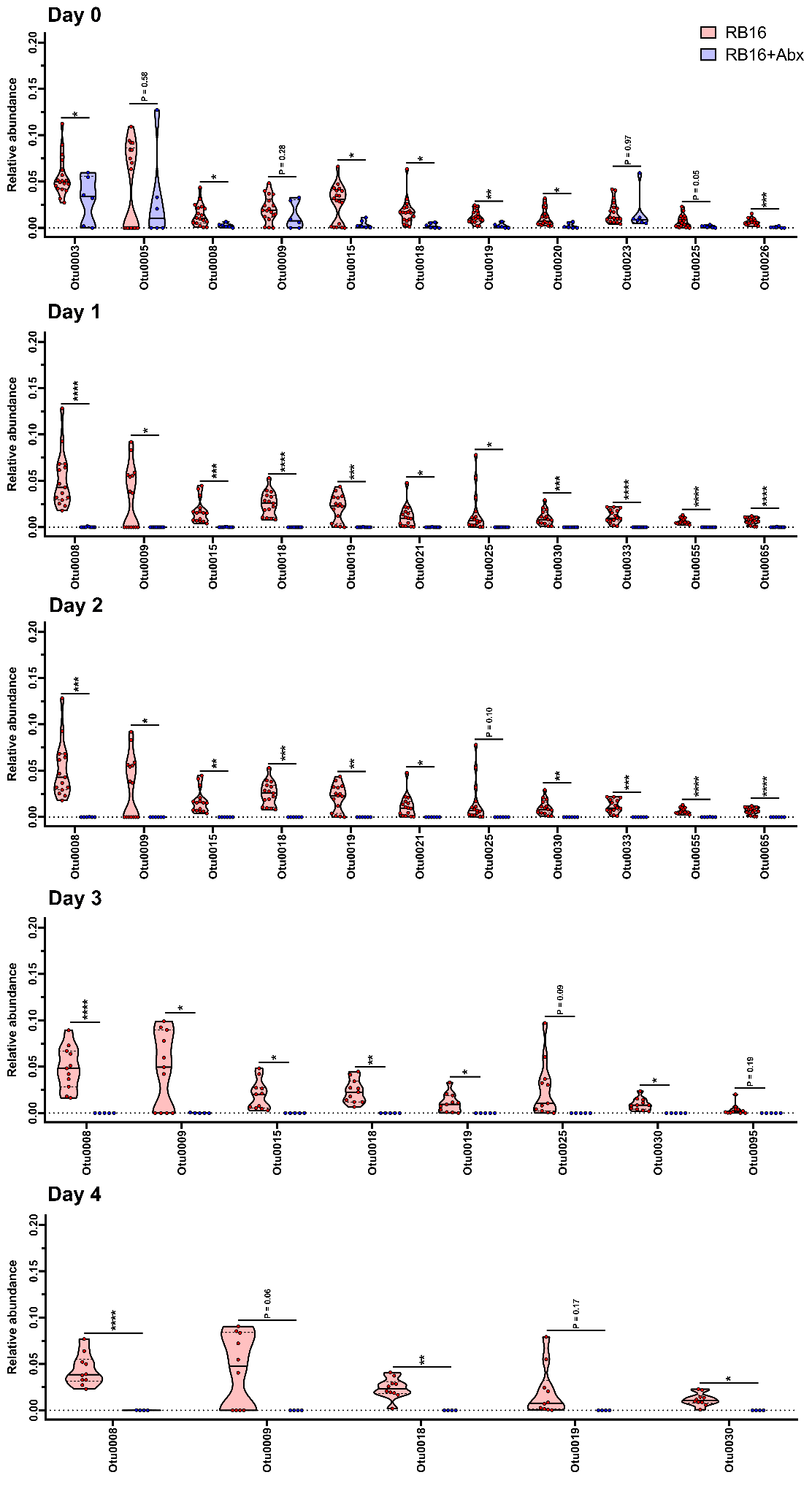

**Supplementary Fig. 16: OTUs that differentiate the microbiota of RB16 from RB07 are sensitive to antibiotic treatment.**

Fecal pellets collected daily from male and female C57BL6/J mice sourced from barriers RB16 and RB16+Abx (n = 10-18 mice per group) following Kp inoculation were subjected to 16S rRNA gene sequencing. Relative abundance of specific OTUs that were differentially abundant between the fecal microbiota of RB16 and RB07 by LEfSe are displayed (*P < 0.05, **P < 0.005, ***P < 0.0005, ****P < 0.00005, Student’s *t* test).

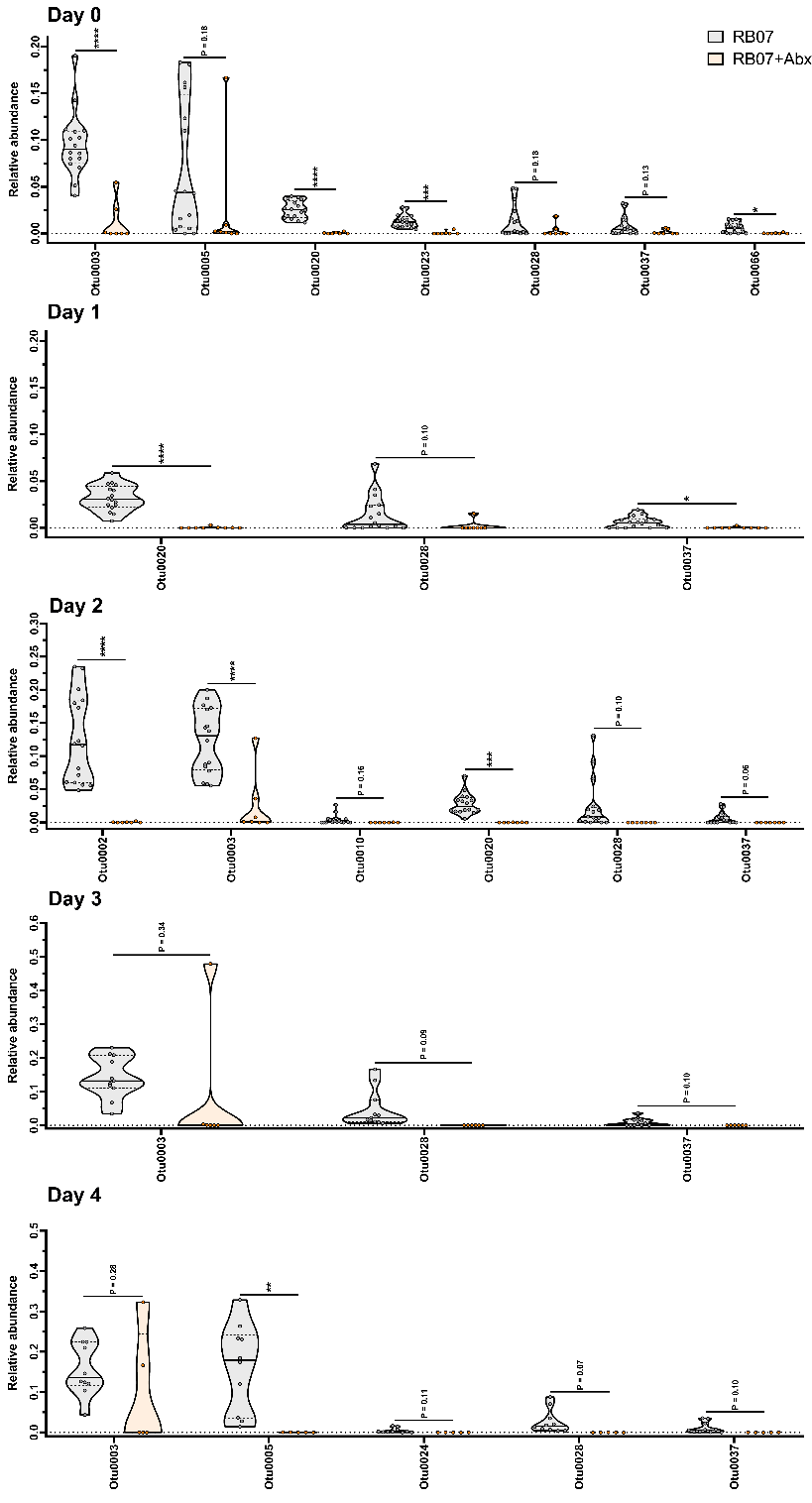

**Supplementary Fig. 17: OTUs that differentiate the microbiota of RB07 from RB16 are sensitive to antibiotic treatment.**

Fecal pellets collected daily from male and female C57BL6/J mice sourced from barriers RB07 and RB07+Abx (n = 9-16 mice per group) following Kp inoculation were subjected to 16S rRNA gene sequencing. Relative abundance of specific OTUs that were differentially abundant between the fecal microbiota of RB16 and RB07 by LEfSe are displayed (*P < 0.05, **P < 0.005, ***P < 0.0005, ****P < 0.00005, Student’s *t* test).

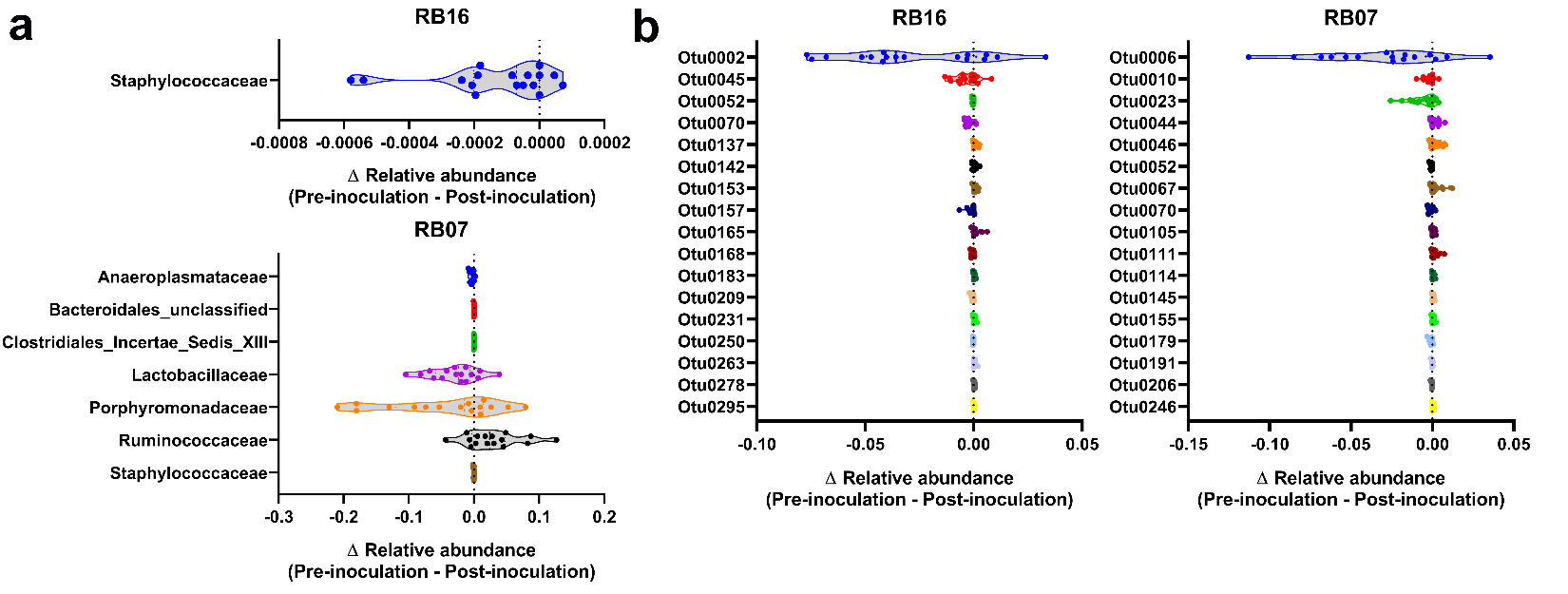
 **Supplementary Fig. 18: Impact of Kp inoculation on the stability of microbiota of RB16 and RB07.**

**a**,**b**, Family (**a**) and OTU (**b**) relative abundance values pre- (Day 0) and post-Kp inoculation (Day 1) were subtracted to determine the impact of Kp inoculation on the fecal microbiota communities of barriers RB16 and RB07. Only significant differential relative abundance values are displayed (median and IQR displayed, one-sample t test compared to a hypothetical value of 0). Each data point represents an individual animal.
